# Deciphering context-dependent epigenetic program by network-based prediction of clustered open regulatory elements from single-cell chromatin accessibility

**DOI:** 10.64898/2026.03.17.712366

**Authors:** Seonjun Park, Sungkook Ma, Wonhyo Lee, Sung Ho Park

**Affiliations:** Department of Biological Sciences, Ulsan National Institute of Science & Technology (UNIST), Ulsan, Korea

## Abstract

Large cis-regulatory domains, spanning tens to hundreds of kilobases, are pivotal in orchestrating cell-state-specific transcriptional programs that define cellular identity. However, existing single-cell analytical frameworks lack the capacity to identify these higher-order structures, thereby obscuring the coordinated, domain-level epigenetic regulation essential for complex biological processes. To address this, we introduce enCORE, a computational framework that leverages enhancer-enhancer interaction networks to determine Clustered Open Regulatory Elements (COREs) solely from single-cell ATAC-sequencing data. Our approach faithfully recapitulates established hematopoietic hierarchies and resolves lineage-specific regulatory programs by recovering canonical master transcription factors, frequent chromatin interactions, and enrichment of fine-mapped immune-related disease-associated genome-wide association study (GWAS) variants. In colorectal cancer, enCORE captures tumor-associated H3K27ac landscapes and prioritizes *USP7* as a potential therapeutic candidate, supported by *in silico* perturbation. Collectively, our framework provides a powerful and scalable platform for deciphering the complex epigenetic architectures underlying human development and disease.

## Main

Gene expression programs are orchestrated not only by discrete regulatory elements, but also by complex interactions within large cis-regulatory domains. Spanning tens to hundreds of kilobases, these domains serve as integrative hubs that consolidate multiple inputs to ensure robust and precise transcriptional control. Their architectures are remarkably diverse, encompassing both activating ensembles, such as super-enhancers (SEs)^1,2^, stretch enhancers^3^, and broad H3K4me3 domains^4^, and repressive domains, including super-silencers^5,6^ and Lamina-associated domains^7^. These domains constitute the functional backbone of the non-coding genome, mediating cell identity specification, context-dependent transcriptional regulation, and the manifestation of disease-associated phenotypes.

Among these large cis-regulatory domains, highly interactive enhancer clusters, recognized as SEs or highly inter-connected enhancers^8^, are well-established mediators of stable transcriptional activation that orchestrate robust expression of lineage-determining and disease-associated genes. These enhancer clusters are marked by three key characteristics: (1) coordinated chromatin accessibility across broad H3K27ac domains^9–11^, (2) dense transcription factor (TF) occupancy, which facilitates the cooperative assembly of regulatory complexes, and (3) frequent enhancer-enhancer chromatin interactions^8,12^ that sustain spatial proximity to ensure potent transcriptional output. Furthermore, the significant enrichment of genome-wide association studies (GWAS) variants within these regions^2^ provides a critical mechanistic framework for interpreting noncoding disease-risk loci.

A defining attribute of these enhancer clusters is their cellular specificity, with their activity and spatial conformation of such clusters being highly dynamic across cell types, developmental stages, and disease states. Traditionally, these domains have been characterized using bulk-level assays, such as ChIP-seq, which inevitably yield averaged signals that obscure cell-specific regulatory nuances. Therefore, resolving these domains at single-cell resolution is imperative to stratify heterogeneous populations and decode the discrete regulatory programs governing development and disease.

The advent of single-cell ATAC sequencing (scATAC-seq)^13^ has enabled the interrogation of epigenetic landscapes at single-cell resolution, thereby overcoming the limitations of bulk-level profiling in resolving cellular heterogeneity. scATAC-seq provides comprehensive maps of cis-regulatory elements^14–16^, allowing for the identification of key regulatory elements that govern cell-type-specific functions and disease-associated phenotypes. Importantly, scATAC-seq-based accessibility maps facilitate the functional interpretation of noncoding variants associated with complex human traits^15^. As a robust and scalable platform, scATAC-seq provides an unprecedented opportunity to examine large-scale, highly interactive enhancer clusters at single-cell resolution across diverse cellular contexts.

Despite the profound biological significance of highly interactive enhancer clusters, robust computational frameworks to predict such large cis-regulatory domains solely from scATAC-seq data are still limited. Current standard workflows predominantly operate at the level of discrete peaks, rendering most scATAC-seq analyses inherently element-centric. Consequently, they are poorly suited to capture the coordinated and cooperative behavior of enhancers underlying highly interactive enhancer clusters. Although a few approaches have been proposed to model such domains, their performance remains challenged by the sparsity and noise inherent to single-cell profiles^17,18^. These limitations underscore the need for a novel analytical framework capable of resolving highly interactive enhancer clusters at single-cell resolution, enabling a more comprehensive understanding of cell-state-specific gene regulation.

Here, we develop enCORE, a framework to identify clustered open regulatory elements (COREs) from scATAC-seq data by integrating three hallmark features of highly interactive enhancer clusters: coordinated chromatin accessibility, dense TF occupancy, and frequent chromatin interactions. Applying enCORE to human peripheral blood (PB), we demonstrate that it precisely recapitulates cell-type-specific transcriptional regulation, with COREs showing marked enrichment for enhancer-enhancer interactions and immune-related disease-associated GWAS variants. In the bone marrow (BM), enCORE effectively captures the epigenetic trajectories of hematopoietic lineage commitment, revealing a lineage-matched maturation continuity from BM to PB. Finally, we show that enCORE elucidates epigenetic reprogramming in colorectal cancer (CRC), providing new insight into disease mechanisms and identifying potential therapeutic candidates.

## Results

### Overview of the enCORE framework

We developed enCORE, a computational framework designed to reconstruct highly interactive enhancer clusters directly from scATAC-seq data (Fig. 1a). The core architecture of enCORE identifies Clustered Open Regulatory Elements (COREs) through a multi-step refinement procedures: (1) extracting enhancer candidates and constructing initial enhancer modules; (2) merging two nearest, disconnected enhancer module with similar inferred gene-regulatory relationships (Link loss correction); (3) estimating the genomic distance threshold for valid interactions to partition merged enhancer module (Distance threshold Optimization); and (4) performing additional rescue step for compensating the sparsity (Sparsity correction) (Methods). The framework then constructs a pseudo-directed enhancer-enhancer interaction network, followed by personalized PageRank-based prioritization (Methods; Fig. 1b). enCORE offers two functional options: a ‘potential’ option for COREs discovery, and an ‘active’ option that incorporates promoter accessibility to emphasize active regulatory programs engaged in the current cell state (Methods).

**Fig. 1:**
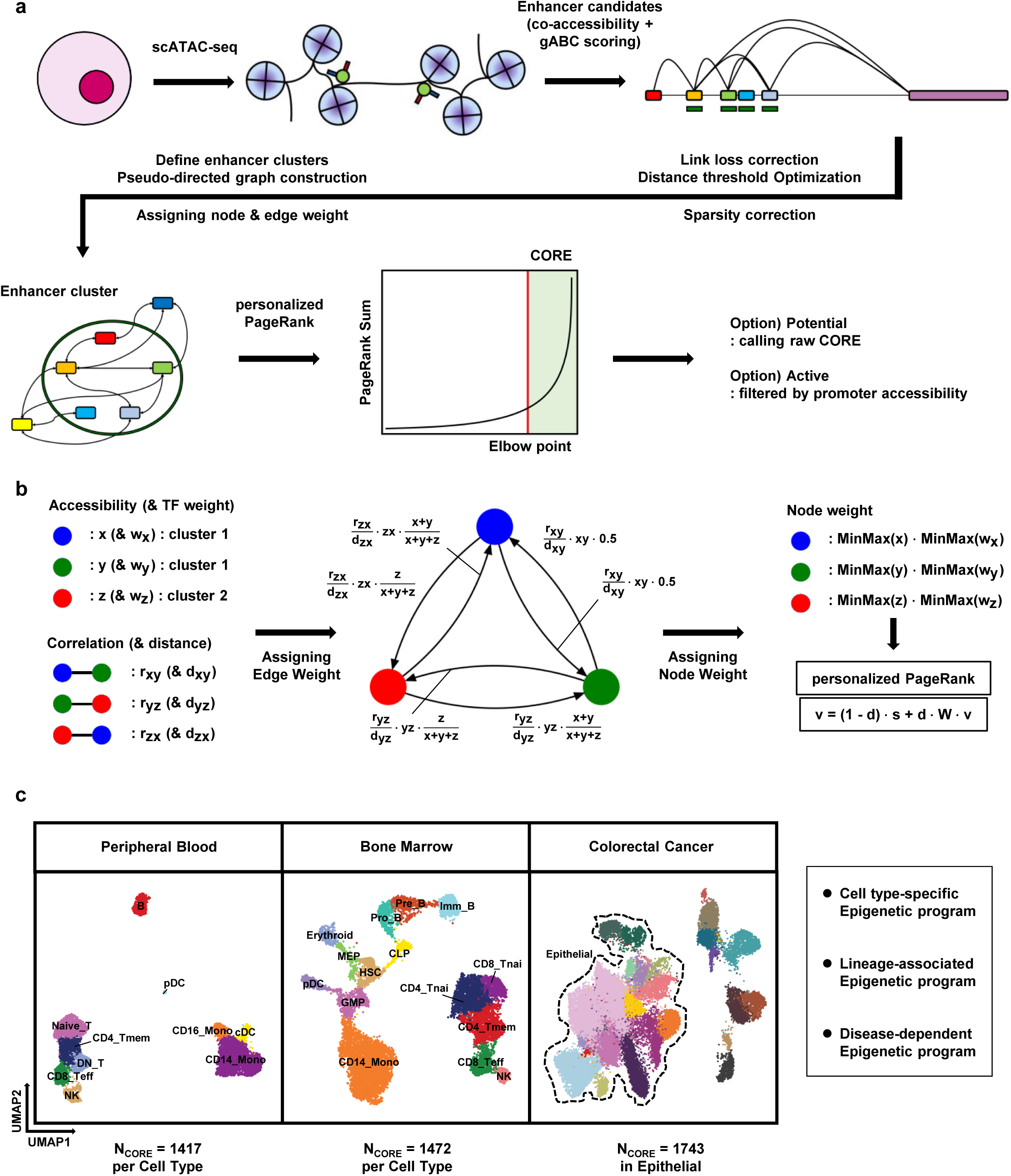
Overview of enCORE and its application to single-cell chromatin accessibility datasets a, The enCORE framework. Step 1: Extracting enhancer candidates using co-accessibility and gABC scoring. Step 2: Link loss correction and sparsity correction with distance threshold optimization. Step 3: Performing pseudo-directed graph construction on defined enhancer clusters. Step 4: Enhancer cluster ranking by personalized PageRank. Step 5: Two options for CORE calling; potential and active. **b,** Detailed procedures for pseudo-directed graph construction. Converting an undirected co-accessibility network between enhancers to a directed graph. Left: Information to be used for network edge and node weight assignment. Middle: Assigning edge weights derived from accessibility of enhancers, co-accessibility coefficient, and contact frequency inferred by the inverse of distance. Right: Assigning node weights derived from the accessibility and TF information of enhancers with MinMax scaling. Node weights were used as the personalization vector for PageRank calculation. **c,** Three single-cell ATAC datasets on which we applied enCORE to extract clustered open regulatory elements (CORE). UMAP embeddings of each dataset, colored and annotated by cell type. The average number of CORE per cell type of each dataset is represented below the UMAP embeddings.

To validate the enCORE framework, we utilized three scATAC-seq datasets: human peripheral blood mononuclear cells (PBMC), bone marrow mononuclear cells (BMMC), and colorectal cancer (CRC) (Methods; Fig. 1c). We extracted COREs from these datasets (Supplementary Table 1-3). We also collected publicly available fine-mapped GWAS variants from CausalDB, curated GWAS variants from Hnisz *et al.*^2^, SEs (in CD4+ T, CD8+ T, NK, B, and CD14+ Monocytes) from H3K27ac bulk ChIP-seq data in SEdb^19^, Hi-C for CD19+ B cells and CD14+ Monocytes, HiChIP for H3K27ac from HiChIPdb, and metastasis-related genes from metsDB and TCGA GDC (Methods).

### COREs define cell type-specific transcriptional programs in PBMC

We first applied enCORE to a 10k PBMC scATAC-seq dataset, identifying COREs across seven immune cell types: CD4+ Memory T cells, CD8+ Effector T cells, B cells, NK cells, CD14+ and CD16+ Monocytes, and classical Dendritic cells (Supplementary Table 1). We hypothesized that COREs serve as primary determinants of cell identity, similar to SEs^2^. To test this, we compared COREs with non-CORE active enhancers (non-CORE), defined as merged SEs and typical enhancers (TEs) regions that do not overlap COREs. Genes associated with COREs exhibited significantly higher cell-type specificity compared with non-CORE-associated genes (Fig. 2a). This trend consistently observed across both enCORE options (Extended Data Fig. 1a,b). The validity of our algorithm in capturing enhancer activity was further corroborated by the substantial overlap between enCORE-inferred enhancers and H3K4me1 peaks in each cell type (Fig. 2b). Notably, COREs were associated with lineage-defining master regulators, such as *GATA3* in NK/T cells, *EBF1* in B cells, and *MAFB* in myeloid cells (Fig. 2c). Publicly available PBMC RNA-seq data revealed that CORE-associated genes display markedly more pronounced cell type-specific transcriptional activation than those associated with non-CORE regions (Fig. 2d, Extended Data Fig. 1c). This transcriptional specificity was also evident in the expression profiles of hematopoietic master regulators (Fig. 2e). TF motif enrichment analysis further confirmed that CORE exhibited stronger enrichment for cell type-specific TF motifs compared with non-CORE regions (Fig. 2f). Overall, these results support the ability of enCORE to recapitulate cell type-specific transcriptional programs in PBMC.

**Fig. 2:**
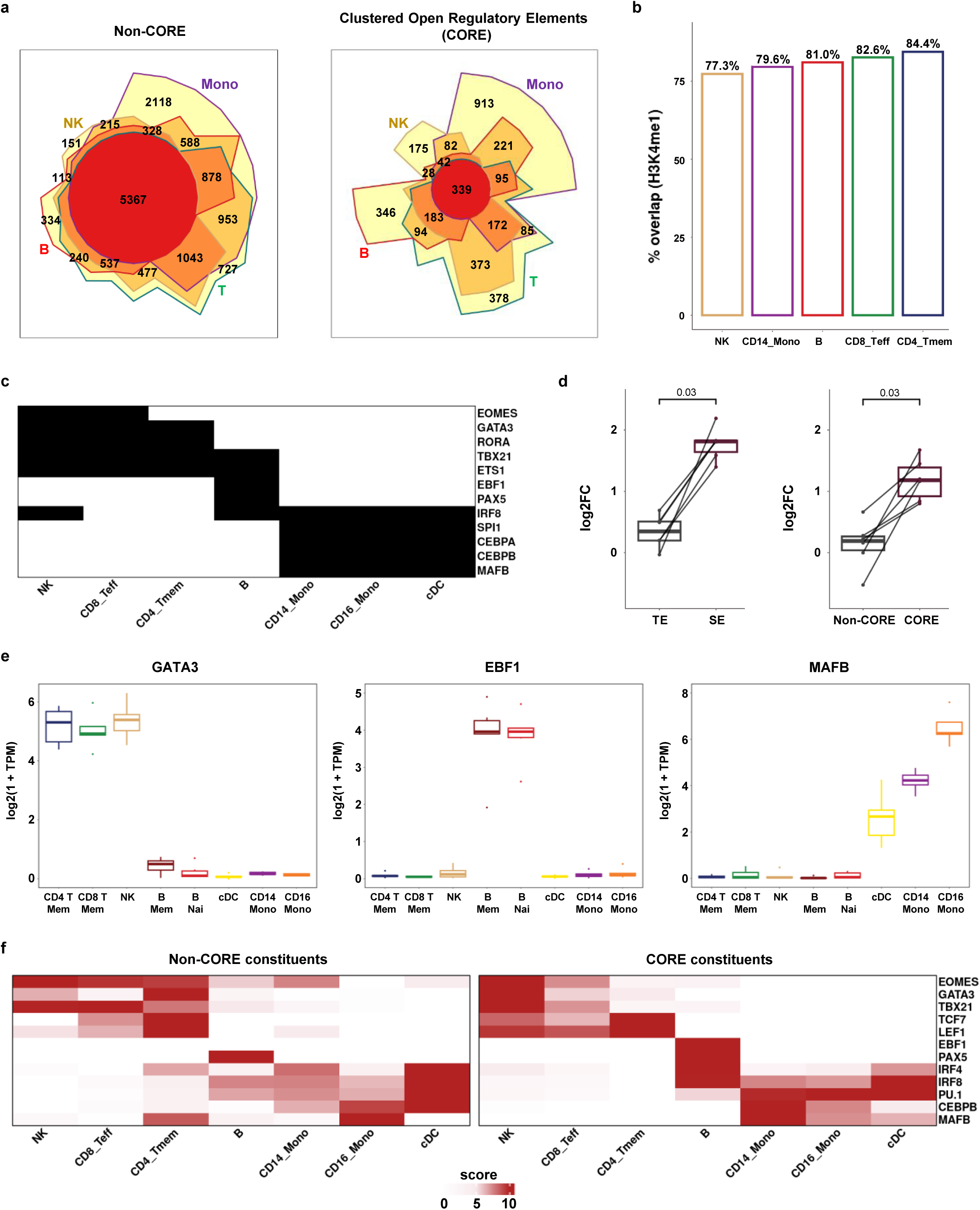
enCORE uncovers cell type-specific transcriptional regulation in PBMC a, Chow-Ruskey diagram of non-CORE-associated genes and CORE-associated genes in PBMC. B cells (red border), Monocytes (purple border), T cells (green border), NK cells (brown border). Monocytes represent the union of CORE-associated genes derived from CD14+ and CD16+ Monocytes, while T cells represent the combined set of CORE-associated genes derived from CD4+ and CD8+ T cells. The color of the borders around each intersection corresponds to the cell types whose genes overlap. The area of each intersection is proportional to the number of genes within the intersection. **b,** Proportion of CORE constituents that overlap with H3K4me1 peaks, relative to the total number of enhancer candidates in each cell type. **c,** Heatmap for cell type-specific master regulators. Black: gene present in the CORE-associated genes of the given cell type. White: absent. **d,** Log2-transformed fold changes of normalized TPM of genes in each cell type. Normalized TPM values were retrieved from the PBMC Monaco dataset in the Human Protein Atlas (HPA). Left: comparison between TE-associated genes and SE-associated genes. Right: comparison between non-CORE-associated genes and CORE-associated genes. P-value was calculated by a two-sided Wilcoxon signed-rank test. **e,** Representative RNA expression for *GATA3*, *EBF1*, and *MAFB*. Normalized TPM values were retrieved from the HPA PBMC dataset. **f,** Enrichment analysis of known TF motifs within CORE and non-CORE constituents across cell types. Colored by enrichment score, defined as modified MinMax-scaled statistical significance (-log10(P-value)). Cell types were indicated as abbreviations (**b**, NK: NK cells, CD14_Mono: CD14+ Monocytes, B: B cells, CD8_Teff: CD8+ Effector T cells, CD4_Tmem: CD4+ Memory T cells; **e**, CD4 T Mem: CD4+ Memory T cells, CD8 T Mem: CD8+ Memory T cells, NK: NK cells, B mem: Memory B cells, B nai: naïve B cells, cDC: classical Dendritic cells, CD14 Mono: CD14+ Monocytes, CD16 Mono: CD16+ Monocytes).

Given that enCORE leverages network structure and TF motifs, rather than the coactivator-driven signal intensity used to define SEs, we reasoned that COREs would capture overlapping yet distinguishable regulatory landscapes. ROC analysis revealed a substantial overlap between COREs and SEs, but also demonstrated that COREs represent a genomic profile distinct from that of SEs (Extended Data Fig. 1d,e and Supplementary Table 4). In monocytes, both COREs and SEs were enriched for cell type-specific GO terms relative to TE, yet they recovered partially distinct genes (Supplementary Fig. 1a,b). For example, in CD14+ Monocytes, COREs uniquely identified key regulators of macrophage polarization and differentiation, such as *KLF4*^20^ and *RBPJ*^21^, whereas bulk SEs exclusively captured the canonical monocyte marker *LYZ*^22^ (Supplementary Fig. 2a-c). These observations suggest that COREs and SEs highlight layers of regulatory features that are shared yet distinct within the same cell population, with COREs prioritizing dynamic regulatory transitions that may be obscured in traditional intensity-based assays.

We next evaluated whether SEs defined from H3K27ac single-cell CUT&Tag-seq (scCUT&Tag-seq)^23^ could detect highly interactive enhancer clusters to an extent comparable to COREs. We identified SEs from a PBMC H3K27ac scCUT&Tag-seq dataset using the ROSE algorithm and benchmarked them against both enCORE-defined COREs and bulk-derived SEs (Methods; Extended Data Fig. 2a,b, Supplementary Table 5). We hypothesized that the inherent sparsity of scCUT&Tag-seq data^23^ might limit the detection of these higher-order architectures. Indeed, while scCUT&Tag-derived SEs showed cell-type-specific distribution patterns, they consistently underperformed in identifying lineage-defining master regulators (Extended Data Fig. 2c,d). Furthermore, although scCUT&Tag-derived SE showed a trend toward higher mRNA-level cell-type specificity than TEs, the difference was not statistically significant (Extended Data Fig. 2e). Collectively, these results underscore the superior practical utility of enCORE; by leveraging network structure and TF motifs, it recovers highly interactive regulatory hubs more effectively than current experimental scCUT&Tag-seq in sparse data regimes.

### COREs capture frequent enhancer-enhancer interactions and poised regulatory states

We next assessed whether COREs reflect the higher-order chromatin topological organization characteristic of interactive enhancer clusters, which is essential for robust, cell state-specific transcription. Interrogation of both H3K27ac HiChIP (GM12878) and Hi-C (CD19+ B cells) datasets revealed a significant enrichment of chromatin interactions within CORE constituents compared with non-CORE regions (Fig. 3a,b). This enrichment was consistently validated across diverse immune lineages, such as CD4+ T cells and CD14+ Monocytes (Extended Data Fig. 3a,b), confirming that COREs preferentially identify chromatin interaction-enriched enhancer architectures across diverse immune cell types.

**Fig. 3:**
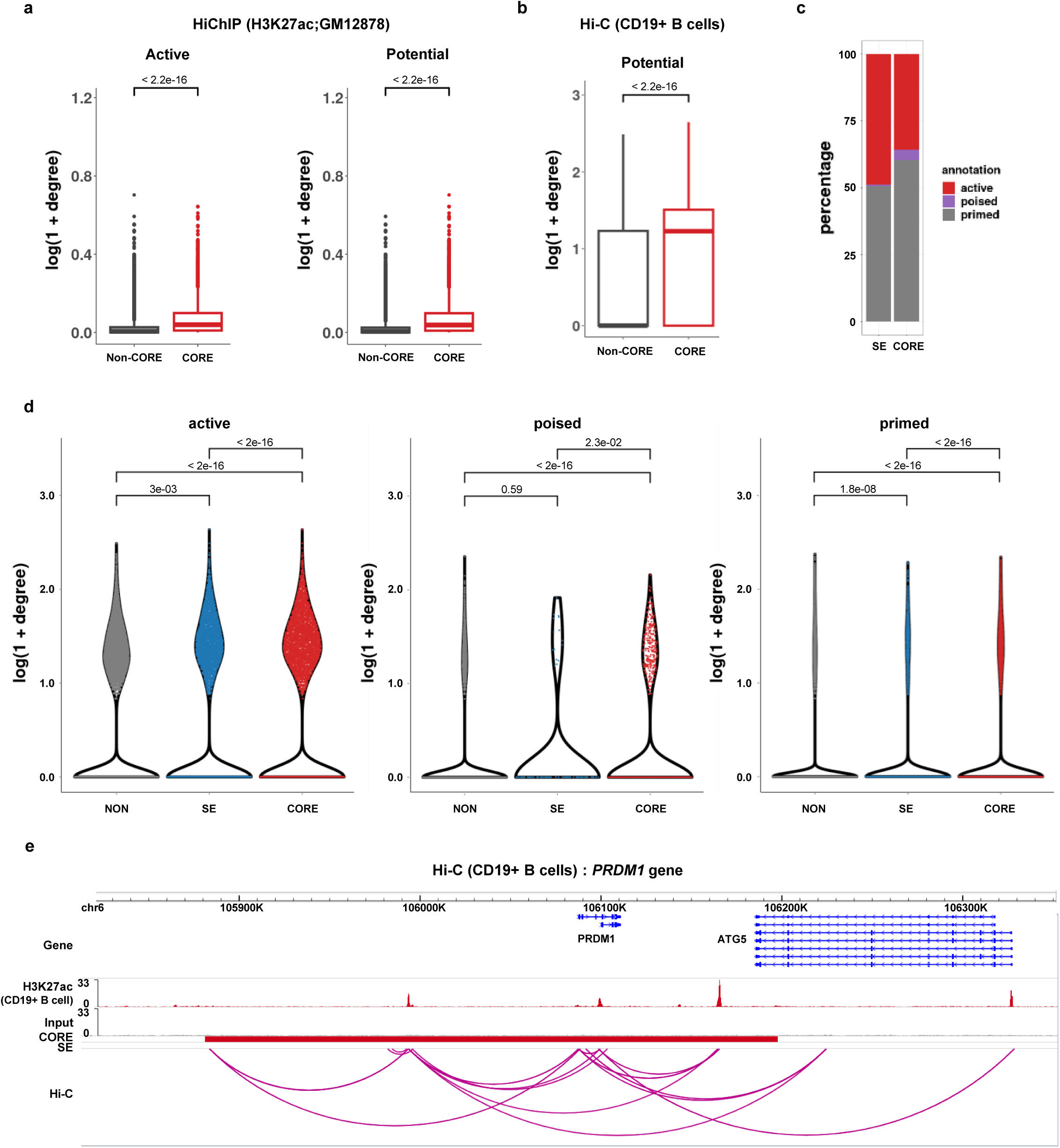
CORE constituents exhibit enriched chromatin interactions a, Log10-transformed weighted degrees of constituents within COREs and non-COREs. Data was retrieved from the H3K27ac HiChIP dataset for GM12878 cells. Left: COREs from the active option, Right: COREs from the potential option. **b,** Log10-transformed weighted degrees of H3K4me1 peaks within COREs and non-COREs (the potential option). Data was retrieved from the Hi-C dataset for CD19+ B cells. Only Hi-C interactions overlapping with H3K4me1 peaks are considered. **c,** Bar graphs showing the relative proportion of each enhancer category from CD19+ B cells within COREs and SEs. active: active enhancers, poised: poised enhancers, primed: primed enhancers that do not fall into either the active or poised categories. **d,** Log10-transformed weighted degrees of H3K4me1 peaks in each enhancer category from CD19+ B cells. Left: active enhancers, Middle: poised enhancers, Right: primed enhancers. Peak group marked by NON represents H3K4me1 peaks that do not fall into either COREs or SEs. **e,** Track visualization at the *PRDM1* locus with Hi-C interactions from CD19+ B cells. P-values were calculated to measure statistical significance (**a,b**, two-sided Wilcoxon rank sum test; **d**, Kruskal-Wallis test with Dunn’s test. P-values were adjusted by the Holm correction method).

Recognizing that enhancer activity can confound comparisons of context-dependent interaction strength^24^, we categorized H3K4me1-marked enhancers within COREs and SEs as active, poised, or primed states based on H3K27ac and H3K27me3 profiles (Methods). Notably, COREs contained a higher proportion of poised enhancers relative to SEs (Fig. 3c). This suggests that while SEs primarily mark current transcriptional output, COREs possess better sensitivity for capturing transcriptionally poised states, potentially representing a reservoir of regulatory potential for future activation. Additionally, we verified that enCORE’s ‘active’ option effectively filters these poised elements, allowing for the precise isolation of regulatory programs actively engaged in the current cell state (Supplementary Fig. 3a,b).

We further compared enhancer-enhancer chromatin interactions within COREs and SEs, stratified by their epigenetic states. In CD19+ B cells, COREs showed consistently higher interaction frequencies across all three enhancer categories relative to SEs (Fig. 3d). Notably, enCORE uniquely identified a higher-order regulatory domain in the vicinity of the *PRDM1* gene, a key regulator of B cell-to-plasma cell differentiation^25,26^, which was characterized by robust intra-domain connectivity that remained undetected by bulk SE assays (Fig. 3e). In CD14+ Monocytes, interaction frequencies within COREs were comparable to those in SEs across most states, with poised COREs showing a marginal, though not statistically significant, increase in the degree of interaction (Extended Data Fig. 3c). Collectively, these results demonstrate that enCORE is a highly effective platform for leveraging complex chromatin topologies, robustly capturing highly interactive enhancer clusters directly from sparse single-cell data.

### Enrichment of immune-related disease-associated GWAS variants within COREs

We next investigated whether fine-mapped GWAS variants are preferentially localized within COREs. Using a fold-change-based enrichment score relative to the genomic background (Methods), we observed that both COREs and SEs exhibit a pronounced enrichment of variants associated with immune-related diseases compared with non-immune traits (Fig. 4a, Extended Data Fig. 4a, Supplementary Fig. 4a). Notably, this enrichment pattern aligned with established disease-critical cell types^27–30^. For instance, fine-mapped variants for Chronic Lymphocytic Leukemia (CLL) and Systemic Lupus Erythematosus (SLE) were enriched within B cell-derived COREs. In contrast, variants for Asthma and Type 1 Diabetes (T1D) were predominantly found within T cell-derived COREs (Fig. 4b,c, Extended Data Fig. 4b-d, Supplementary Fig. 4b,c). Interestingly, in Multiple Sclerosis (MS), we observed a striking enrichment of risk variants within CD8+ T-cell and B-cell COREs (Extended Fig. 4e, Supplementary Fig. 4d), corroborating recent shifts in our understanding of MS pathogenesis beyond traditional CD4+ T-cell-centric models.^31–33^. These cell-type-resolved enrichments were validated across multiple GWAS cohorts (Extended Data Fig. 4, Supplementary Fig. 4). Furthermore, motivated by the lipid-response signature observed in monocyte COREs (Supplementary Fig. 1a,b), we examined coronary artery disease (CAD) risk across three independent cohorts. CAD risk loci showed the strongest enrichment in CD14+ Monocyte COREs (Extended Data Fig. 5a-d). A representative example is the *LIPA* locus, where a disease-relevant enhancer cluster was uniquely resolved by enCORE (Extended Data Fig. 5e). Given the pivotal role of *LIPA* in lipid metabolism^34^ and its reported eQTL association in monocytes^35^, our findings implicate CORE-mediated regulation of *LIPA* as a key candidate mechanism underlying CAD risk.

**Fig. 4:**
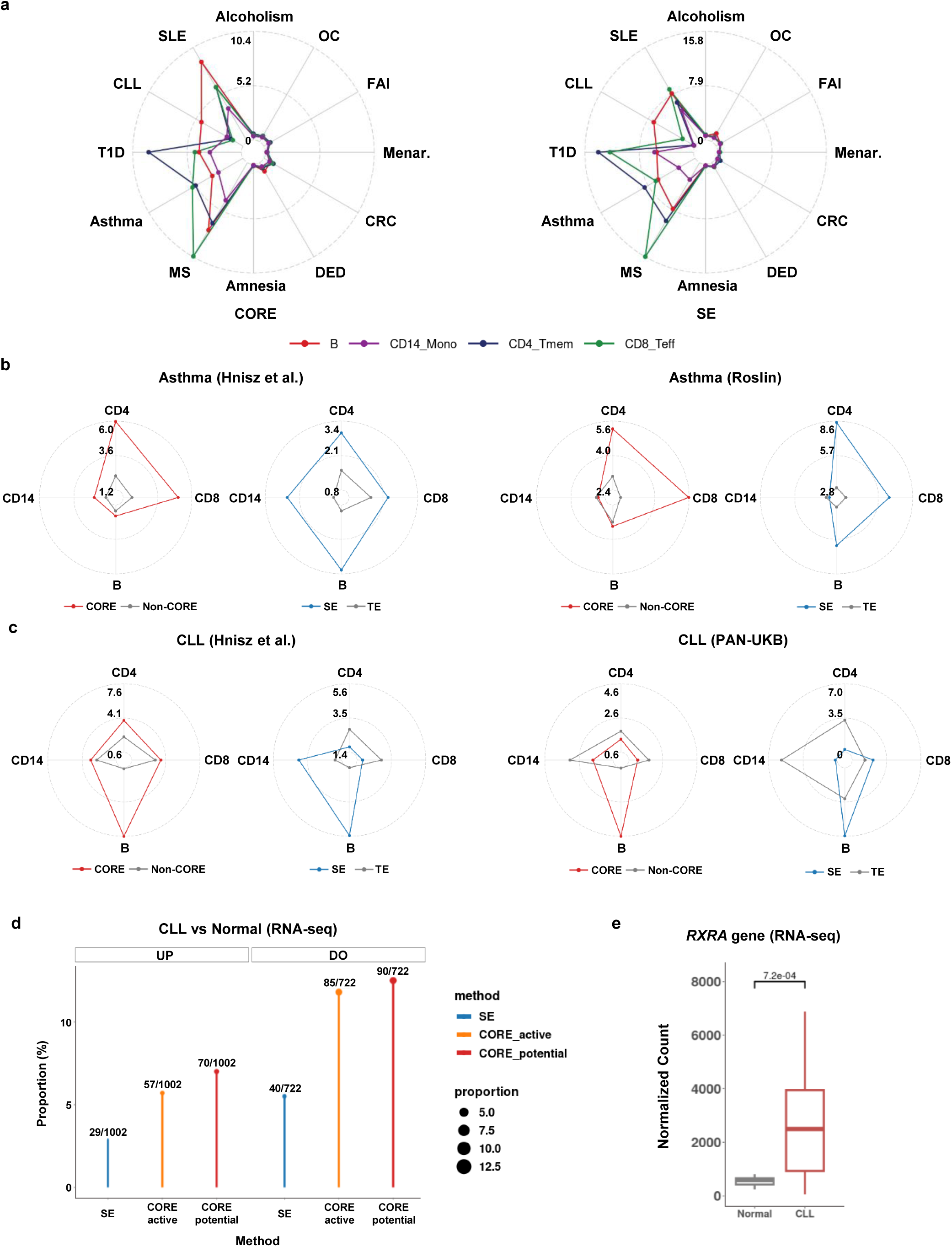
Fine-mapped GWAS variants for immune-related diseases are enriched in COREs a, Enrichment analysis of fine-mapped GWAS variants for immune-related diseases or non-immune-related traits in COREs (left) and SEs (right). SLE: Systemic Lupus Erythematosus, CLL: Chronic Lymphocytic Leukemia, T1D: Type 1 Diabetes, Asthma: Asthma, MS: Multiple Sclerosis, Amnesia: Amnesia, DED: Dry Eye Disease, CRC: Colorectal Cancer, Menar.: Menarche, FAI: Fasting Insulin, OC: Ovarian Cancer, Alcoholism: Alcoholism. **b-c,** Enrichment analysis of Young Lab-curated (Hnisz *et al.*) and fine-mapped GWAS variants for **b,** Asthma; **c,** CLL in COREs and SEs. Lines are colored by region categories: COREs (red color), SEs (blue color), and TEs (grey color). B: B cells, CD14: CD14+ Monocytes, CD4: CD4+ Memory T cells, CD8: CD8+ Effector T cells. **d,** Proportion of differentially expressed genes between CLL and Normal B cells overlapping CORE- or SE-associated genes. UP: up-regulated genes in CLL, DO: down-regulated genes in CLL. CORE_potential: COREs from the potential option (red color), CORE_active: COREs from the active option (orange color), SEs (blue color). **e,** Box plot showing a comparison of the *RXRA* gene expression between CLL and Normal B cells. Read counts were normalized by the Relative Log Expression (RLE) method. P-value was calculated using the Wald test in DESeq2.

Having established the B cell-specific enrichment of CLL risk variants (Fig. 4c), we next assessed the sensitivity of enCORE in capturing CLL-associated transcriptional changes following disease onset. COREs identified a broader repertoire of differentially expressed genes than bulk-derived SEs, with the ‘potential’ option providing the highest recovery of disease-relevant signatures (Fig. 4d). Importantly, the *RXRA* locus was identified exclusively by enCORE (Fig. 4e). Given that *RXRA* expression is markedly elevated in both human and murine B-CLL cells^36^, our findings suggest that the domain-level epigenetic remodeling of this locus may play a pivotal role in CLL pathogenesis. In summary, enCORE provides a cell-type-resolved framework to interpret fine-mapped GWAS variants by linking non-coding disease risk loci to domain-scale regulation of disease-associated genes.

### enCORE reconstructs hematopoietic trajectories and lineage-defining regulators in human BMMC

To determine if CORE architectures delineate the logic of hematopoietic lineage commitment, we applied enCORE to a human BMMC scATAC-seq dataset (Supplementary Table 2). By quantifying the cosine similarity of CORE profiles, we reconstructed the epigenetic proximity governing the hematopoietic hierarchy (Supplementary Table 6a). These patterns faithfully recapitulated the established hematopoietic lineage trajectory (Fig. 5a, Extended Data Fig. 6a,b). We observed a notable epigenetic association among Granulocyte-Myeloid-Progenitors (GMP), Common Lymphoid-Progenitors (CLP), and plasmacytoid Dendritic Cells (pDC) in the potential option, supporting reports that pDC can originate from both lymphoid and myeloid lineages^37^.

**Fig. 5:**
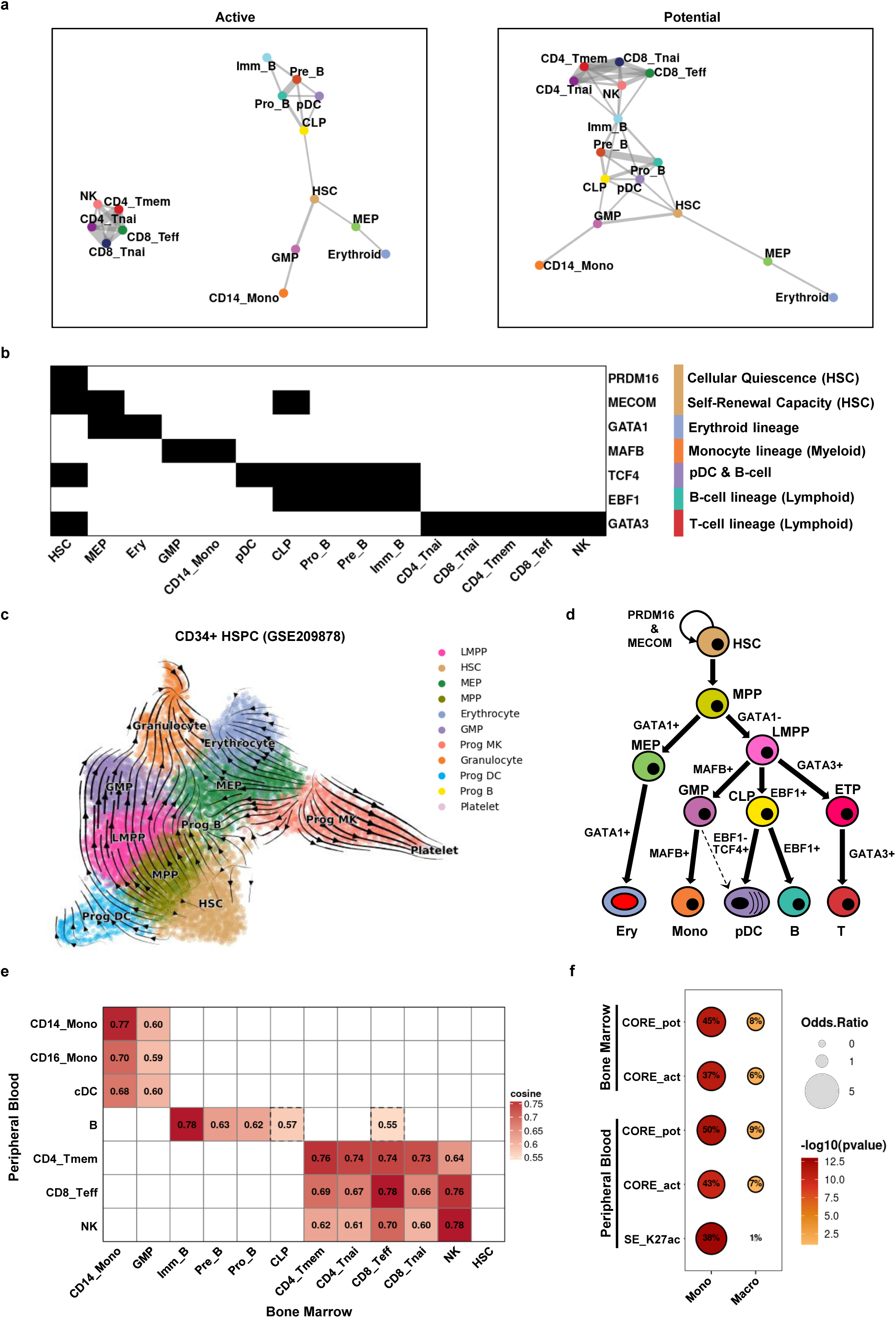
enCORE deciphers lineage-committing regulatory determinants in bone marrow hematopoiesis a, Graph-based representation of the similarity among cell type-specific CORE sets identified in BMMC. Left: COREs from the active option. Right: COREs from the potential option. Nodes are colored and annotated by cell types. Edges are shown only for node pairs with cosine similarity greater than 0.55. **b,** Heatmap for lineage-committing master regulators. Black: gene present in the CORE-associated genes of the given cell type. White: absent. **c,** UMAP coordinates with stream plot of velocity vectors calculated by MultiVelo in CD34+ hematopoietic cells, colored and annotated by cell types. **d,** Model of bone marrow hematopoiesis. **e,** Heatmap for cosine similarity metrics between CORE profiles from bone marrow (BM) and peripheral blood (PB). Red: cosine similarity > 0.55. White: cosine similarity < 0.55. Values in each heatmap cell represent the cosine similarity using the potential option of enCORE. Heatmap cells with dotted borders indicate cases where the cosine similarity < 0.55 using the active option of enCORE. **f,** Dot plot for the enrichment of monocyte-elevated or macrophage-elevated H3K27ac peaks within COREs and SEs. Values shown in each dot represent the percentage of monocyte-elevated or macrophage-elevated H3K27ac peaks that overlap COREs or SEs. Dot size is determined by the common odds ratio. Red/Orange: p-value < 0.05. Grey: p-value > 0.05. Mono: monocyte-elevated H3K27ac peaks. Macro: macrophage-elevated H3K27ac peaks. CORE_pot: COREs from CD14+ Monocytes using the potential option. CORE_act: COREs from CD14+ Monocytes using the active option. SE_K27ac: SEs from CD14+ Monocytes. Cell types were indicated as abbreviations (**a-e**, HSC: Hematopoietic Stem Cells, MPP: Multipotent Progenitors, LMPP: Lymphoid-Myeloid Primed Progenitors, MEP: Megakaryocyte-Erythroid Progenitors, Prog MK: Megakaryocyte Progenitors, GMP: Granulocyte-Myeloid Progenitors, Prog DC: Dendritic cell Progenitors, CLP: Common Lymphoid Progenitors, Prog B: B cell Progenitors, ETP: Early T-cell Precursors, Erythroid or Ery: Erythroid, Erythrocytes: Erythrocytes, Platelet: Platelets, Granulocyte: Granulocytes, Mono: Monocytes, CD14_Mono: CD14+ Monocytes, CD16_Mono: CD16+ Monocytes, cDC: classical Dendritic cells, pDC: plasmacytoid Dendritic cells, B: B cells, Pro_B: Pro-B cells, Pre_B: Pre-B cells, Imm_B: Immature B cells, NK: NK cells, T: T cells, CD8_Teff: CD8+ Effector T cells, CD8_Tnai: CD8+ naïve T cells, CD4_Tmem: CD4+ Memory T cells, CD4_Tnai: CD4+ naïve T cells).

Consistent with the lineage-specific chromatin landscapes, CORE-associated genes effectively recovered key lineage-defining regulators, including *EBF1* (B cell), *GATA3* (NK/T cell), *MAFB* (Monocyte/Myeloid), and *GATA1* (Megakaryocyte/Erythroid) in their respective lineage (Fig. 5b). Within Hematopoietic Stem Cells (HSC), COREs were explicitly identified at the *MECOM* (*EVI1*) and *PRDM16* loci, which are crucial factors for self-renewal^38^ and quiescence^39^, respectively. Interestingly, *MECOM-*associated CORE were also detected in both Megakaryocyte-Erythroid-Progenitors (MEP) and CLP, providing a regulatory basis for reports linking *MECOM* mutations to amegakaryocytic thrombocytopenia and B cell deficiency^40^. These observations underscore the utility of enCORE in linking lineage-resolved regulatory programs to hematopoietic phenotypes.

While lymphoid and myeloid lineages were clearly separated, CLP and GMP shared similar CORE profiles under the potential option. Given that the GMP in our dataset are *GATA1*-negative, our findings support a previously proposed model^41^ where these progenitors are derived from Lymphoid-Myeloid-Primed-Progenitors (LMPP) and share a regulatory trajectory with the lymphoid lineage. To validate this transition, we applied MultiVelo to CD34+ hematopoietic cells (Methods; Fig. 5c), which confirmed that GMP primarily originate from the LMPP pool in this context. Based on these collective findings, we present a schematic representation of bone marrow hematopoiesis (Fig. 5d).

We further asked whether CORE profiles preserve maturation continuity across spatial compartments, specifically from the bone marrow (BM) to peripheral blood (PB). Comparing corresponding lineages between BM and PB, we found that PB CORE profiles most closely matched their late-stage counterparts, with within-lineage ordering well preserved (Fig. 5e and Supplementary Table 6b). For example, PB B cell COREs exhibited the highest similarity to BM immature B cells, with decreasing similarity to Pre-B, Pro-B, and CLP states.

The observed preservation suggested that downstream fates might be epigenetically imprinted earlier in the hematopoietic hierarchy. We therefore hypothesized that BM CD14+ monocyte COREs harbor anticipatory macrophage-biased signatures. Indeed, when compared with macrophage-versus-monocyte differential H3K27ac peaks, both COREs and SEs overlapped with monocyte-elevated peaks, but only COREs showed significant enrichment for macrophage-elevated peaks (Methods; Fig. 5f and Supplementary Table 7).

Genome browser inspection highlighted representative loci, including *DCSTAMP* (osteoclast multinucleation^42^ & macrophage migration^43^) and *MEF2A* (IFN I induction in macrophages^44^), where macrophage-elevated peaks exclusively overlapped with BM- and PB-derived COREs but not SEs (Extended Data Fig. 7 and Supplementary Table 7). Furthermore, macrophage-elevated regulatory elements at the *BHLHE41* locus, a key regulator of monocyte-to-alveolar macrophage differentiation^45^ and macrophage-specific H3K27ac profile^46^, were uniquely recovered within COREs (Methods). Together, these results demonstrate that enCORE retains lineage-biased epigenetic signals that are attenuated in traditional SE-based summaries, providing clues to identify early regulatory programs primed for specific developmental fates.

### enCORE identifies CRC-specific epigenetic reprogramming and therapeutic target candidates

Finally, we evaluated the clinical utility of enCORE in dissecting disease-dependent epigenetic landscapes using a colorectal cancer (CRC) dataset. We extracted COREs from CRC epithelial cells (Supplementary Fig. 5 and Supplementary Table 3) and compared them with SEs derived from two independent bulk H3K27ac ChIP-seq datasets (Methods; Supplementary Table 8-10). To assess disease-state classification, we performed NMF/K-means clustering using H3K27ac profiles from CRC and normal samples (Methods). Notably, CORE-based models distinguished CRC from the normal state, whereas SEs explained only a small fraction of the disease-specific signatures (Fig. 6a, Supplementary Fig. 6a,b, Supplementary Fig. 7a,b,g). This performance was further cross-validated by XGBoost classification (Supplementary Fig. 6c-f, Supplementary Fig. 7c-f), where COREs showed higher log2 fold changes than SEs or TEs (Fig. 6b, Supplementary Fig. 7h). In addition, *de novo* TF motif enrichment analysis revealed that the *FOXM1* TF motif was exclusively enriched within CORE constituents (Fig. 6c). Given the established role of *FOXM1* as a central driver of CRC pathogenesis^47,48^, our results suggest that COREs may serve as convergent regulatory hubs where cancer-specific TF coordinate higher-order chromatin landscapes to drive malignant phenotypes.

**Fig. 6:**
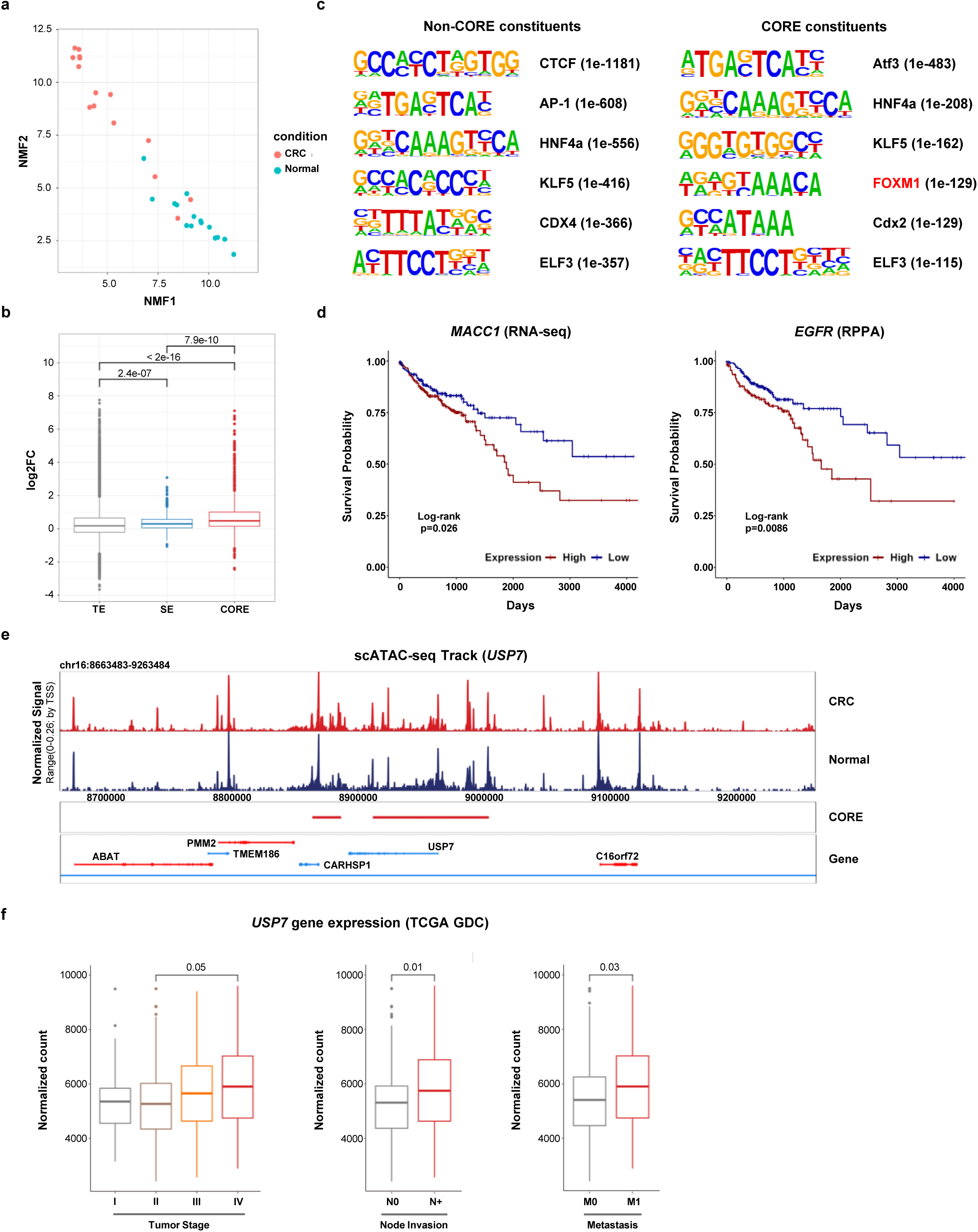
enCORE identifies CRC-associated epigenetic reprogramming and actionable therapeutic targets. a, NMF analysis of H3K27ac ChIP-seq signals within COREs to distinguish between CRC and Normal samples. Dots are colored by disease state: CRC (red color), Normal (blue color). **b,** Log2-transformed fold changes of H3K27ac ChIP-seq signals between CRC and Normal within COREs, SEs, and TEs. Dots and borderlines are colored by region categories: COREs (red color), SEs (blue color), TEs (grey color). **c,** Enrichment analysis of *de novo* TF motifs in CORE and non-CORE constituents. Left: TF motif enrichment analysis using non-CORE constituents, Right: TF motif enrichment analysis using CORE constituents. The *FOXM1* motif is highlighted in red color. **d,** TCGA survival analysis for *MACC1* and *EGFR* genes. Lines are colored by expression level status: High gene expression (red color), Low gene expression (blue color). Survival curves for the *MACC1* and *EGFR* genes were obtained from RNA-seq and RPPA data, respectively. **e,** Track visualization at the *USP7* locus with scATAC-seq signals normalized by reads in TSS. Colored and annotated by disease state: CRC (Colorectal cancer epithelial cells; red color), Normal (Normal epithelial cells; blue color). **f,** Box plots showing *USP7* expression in TCGA GDC RNA-seq data across tumor stage (I-IV; left); lymph node invasion status (N0, no lymph node invasion; N+, lymph node invasion; middle); distant metastasis status (M0, no metastasis; M1, metastasis; right). Read counts were normalized by the Relative Log Expression (RLE) method. P-values were calculated to measure statistical significance (**b**, Kruskal-Wallis test with Dunn’s test. P-values were adjusted by the Holm correction method; **d**, Log-rank test; **f**, Left: Kruskal-Wallis test with Dunn’s test. P-values were adjusted by the Holm correction method. Middle: two-sided Wilcoxon rank sum test. Right: two-sided Wilcoxon rank sum test).

Next, we investigated whether well-known CRC-related master TF candidates were represented within CORE- or SE-associated genes (Supplementary Table 11). While both domains encompassed these candidates (Extended Data Fig. 8), TCGA survival analysis showed significant associations for CORE-associated genes, such as *EGFR*^49^, and *MACC1*^50,51^, were significantly linked to poor clinical outcomes (Fig. 6d). Interestingly, enCORE uniquely identified several genes recently implicated in CRC prognosis, radioresistance, and metastasis^52–54^, including *ISM1*, *PRDM15*, and *HOXC10*, that were omitted by SE-based analyses (Supplementary Fig. 8).

Since three-dimensional (3D) chromatin architecture can potentiate invasion-associated transcriptional programs^55^, we next examined the network topology of metastasis-associated elements. Metastasis-related genes were more abundantly represented within COREs than SEs (Extended Data Fig. 9a,b and Supplementary Table 12,13a). Intriguingly, whereas SEs were characterized by higher node degrees, COREs retained high degrees while exhibiting significantly higher clustering coefficients (Extended Data Fig. 9c). Indeed, enhancers associated with metastasis-related genes showed markedly higher degrees and clustering coefficients than those linked to other genes (Extended Data Fig. 9d,e). This concordance with the network properties of COREs may account for the enhanced recovery of metastasis-related genes within COREs. Beyond CRC, this trend was corroborated in PBMC Hi-C networks, where COREs displayed node degrees comparable to those of SEs but consistently higher clustering coefficients (Extended Data Fig. 9f). Collectively, our results suggest that the locally clustered connectivity captured by enCORE is a critical determinant of invasion-associated transcriptional programs, offering a more nuanced view of the cancer genome than traditional hub-centric models.

We next explored the therapeutic potential of CORE-associated regulatory programs in CRC. Motivated by the exclusive enrichment of the *FOXM1* motif within COREs, we prioritized upstream regulators that could directly regulate *FOXM1*. As a prime candidate, enCORE exclusively identified an enhancer cluster at the *USP7* locus, a critical deubiquitinase that stabilizes *FOXM1*^56^ (Fig. 6e, Extended Data Fig. 10a and Supplementary Table 18). Notably, *USP7* expression increases with advanced tumor stage, lymph node invasion, and metastasis (Fig. 6f), suggesting a major role in CRC progression. Given the emergence of *USP7* as a promising therapeutic target in CRC^57–59^, we performed a computational perturbation experiment using AlphaGenome to test the functional dependency of this USP7-proximal CORE (Methods; Extended Data Fig. 10b,c and Supplementary Table 14). *In silico* knockout of an H3K27ac-marked enhancer peak within the USP7-proximal CORE predicted a significant reduction in *USP7* transcript abundance. This regulatory link was further corroborated by H3K27ac HiChIP data from HT29 cells, which revealed cis-regulatory chromatin loops tethering the USP7-proximal CORE to its cognate promoter (Methods; Extended Data Fig. 10a). These results indicate that the USP7-proximal CORE is a major contributor to *USP7* dysregulation in CRC. In conclusion, our study demonstrates that enCORE is a powerful and scalable platform for elucidating disease-dependent epigenetic programs and prioritizing novel therapeutic interventions.

## Discussion

This study introduces enCORE, a network-based framework that infers highly interactive enhancer clusters from scATAC-seq data by leveraging enhancer-enhancer interaction networks. Unlike traditional workflows, enCORE yields a domain-scale representation of regulatory architecture, integrating coordinated chromatin accessibility, TF occupancy, and network topology. Across diverse biological contexts, enCORE effectively bridges the gap between chromatin organization, cell identity and disease pathogenesis at the single-cell level.

The advent of single-cell epigenomics has exposed the inherent limitations of population-averaged regulatory domains derived from bulk-level profiles, which often blur cell-state heterogeneity and mask subpopulation-specific epigenetic programs. This underscores the necessity of interrogating large cis-regulatory domains directly at single-cell resolution, where domain-scale organization can be resolved in a cell-state-aware manner. enCORE addresses this critical gap by dissecting heterogeneous populations and summarizing context-dependent architectures into highly interactive enhancer clusters. In PBMC and BMMC, for instance, CORE profiles faithfully recapitulated hematopoietic lineage hierarchies and recovered canonical master regulators, supporting that domain-scale summaries remain highly interpretable despite the technical sparsity of single-cell data. Consistent with this, fetal lung analyses showed that enCORE more consistently recovered cell-type-specific master transcription factors than scATAC-derived SE annotations (dbscATAC^18^) or network-centric unsupervised method (eNet^60^), underscoring its robustness to the inherent sparsity of single-cell data (Supplementary Fig. 9 and Supplementary Table 15,16). As atlas-level single-cell datasets continue to accumulate, domain-level representations will become indispensable for investigating state-resolved epigenetic programs.

From the perspective of large cis-regulatory domains, the defining feature of highly interactive enhancer clusters is not any single element, but the cooperative organization of multiple enhancers into a coordinated regulatory assembly stabilized by spatial proximity. enCORE encodes this domain concept by prioritizing enhancer ensembles that exhibit local topological coherence in the enhancer-enhancer interaction networks, effectively transcending element-centric signals. This conceptualization implies that large enhancer domains can be described not only by broad activity marks, but also by network topology that reflects how constituent elements are wired together. In this context, CORE provides a topology-aware domain representation that enables tractable interrogation of how regulatory assemblies are maintained or rewired during cell state transitions at the single-cell level.

While COREs and SEs share a conceptual foundation as enhancer clusters, our results demonstrate that they prioritize distinct facets of regulatory architecture. Importantly, COREs harbor a significantly higher fraction of poised/primed enhancer constituents than SEs, suggesting a unique capacity to capture latent regulatory potential. Moreover, whereas SEs are typically aligned with degree-ranked hubs, COREs favor regions with higher clustering coefficients in the enhancer-enhancer interaction networks. In CRC, COREs preferentially colocalized with cancer-specific H3K27ac programs and prioritized key oncogenic regulators distinct from those identified by SEs, suggesting that enCORE can delineate context-dependent regulatory programs frequently underrepresented in intensity-derived annotations. Furthermore, enCORE captured a high-degree, locally clustered enhancer profile distinct from SEs and eNet, with preferential recovery of metastasis-related programs (Supplementary Fig. 10 and Supplementary Table 13b,16-18). Together, these findings position enCORE as a powerful, complementary strategy to SE for interpreting context-dependent epigenetic regulation at the single-cell level.

Despite its utility, our work has several limitations. While enCORE integrates three primary hallmarks of highly interactive enhancer clusters, the current framework does not explicitly disentangle the relative contribution of each feature to specific CORE assignments across different biological contexts. To address this interpretability gap, future work will incorporate explainable modeling approaches using generalized linear models (GLM) or XGBoost to quantify per-context feature importance and assign per-region attributions. Additionally, while we validated enCORE across diverse human datasets, further investigation is required to assess the cross-species conservation of these higher-order architectures in other organisms, such as mice. Finally, as our evaluations were primarily conducted *in silico*, additional subsequent experimental studies will be essential to establish the definitive causal roles of these domains in gene regulation.

Collectively, enCORE provides a flexible framework for inferring highly interactive enhancer clusters and for decoding domain-level epigenetic programs. By resolving complex regulatory programs from lineage-biased priming to disease-dependent reprogramming, enCORE offers a powerful platform for advancing our understanding of human development and pathology.

## Methods

### enCORE framework

enCORE framework is a workflow based on the following procedures: (1) Enhancer candidates extraction, (2) Initial decomposed components (enhancer clusters) inference, (3) Link loss correction, (4) Automatic distance threshold calculation, (5) Sparsity correction (rescue steps for enhancer pairs with high cosine similarity and lost peaks with high accessibility), (6) Pseudo-directed graph construction (with network node and edge weight assignment) (7) personalized PageRank-based ranking for enhancer clusters, (8) Distilling raw CORE (“potential” option) or filtered CORE (“active” option) after iterative proximal enhancer filtering procedure.

There are two options for analysis with the enCORE framework: potential and active. Because poised enhancers can exhibit high chromatin accessibility^61^ and polycomb domains near lineage-specifying developmental regulators can occur within TADs^62^, the potential option may capture accessible regulatory elements beyond the current transcriptionally active state. To emphasize context-specific activity, we additionally implemented the active option by imposing a promoter-accessibility constraint. More detailed, in the active option, the *iterative proximal enhancer clusters filtering* step conditioned on promoter accessibility was applied to COREs inferred under the potential option.

### gABC scoring & Extracting initial enhancer clusters

Peaks with a co-accessibility (correlation coefficient) of 0.2 or higher were defined as initial enhancer candidates if they were located more than 2kb away from the transcription start site (TSS) of UCSC known genes and showed co-accessibility with at least one promoter. The normalized accessibility (CPM) of these initial enhancer candidates was used as input to the generalized Activity-by-Contact (gABC) scoring procedure implemented in STARE^63^. Peaks with a gABC score greater than 0.01 were redefined as filtered enhancer candidates annotated with gABC scores for multiple genes. For each cell population, we calculated pairwise cosine similarity between co-accessible filtered enhancer candidates based on the gABC score. Subsequently, co-accessibility network edges with a cosine similarity below the specific cut-off (default: 0.4) were removed using the decompose function from the igraph package (v1.5.1). The remaining decomposed components (disconnected clusters in the network) could be defined.

### Link loss correction

We assumed that some co-accessibility network edges may be lost because of the sparsity of scATAC-seq data. This can make more decomposed components compared with the expected number of enhancer clusters. Thus, pairwise cosine similarities between any two components within 500kb were recalculated using the summed values of gABC scores of enhancers in each component to correct the loss of network edges. The farthest two components with a cosine similarity greater than the specific cut-off (default: 0.4) and all intermediate components between them were merged into a single component. The final components obtained after the merging process were defined as Link loss correction (LLS)-corrected enhancer clusters.

### Distance threshold optimization & Sparsity correction

Details of the distance threshold optimization process are as follows. LLS-corrected enhancer clusters are likely to span excessively large genomic distances. Based on the assumption that the likelihood of functionally effective three-dimensional (3D) chromatin interactions decreases with increasing one-dimensional genomic distance, we automatically determined the effective distance threshold for interactions and re-segmented each LLS-corrected enhancer cluster accordingly.

To infer the distance threshold, we employed a strategy that maximizes both the following two conditional probabilities for the genomic distance between the two closest enhancer candidates in the intra-cluster (within a specific enhancer cluster) and inter-cluster (between two enhancer clusters) contexts.

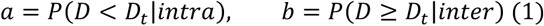

where *P*(*D* < *D*_*t*_|*intra*) is the conditional probability that the intra-cluster distance *D* is smaller than a specific threshold *D*_*t*_, and *P*(*D* ≥ *D*_*t*_|*inter*) is the conditional probability that the inter-cluster distance *D* is bigger than a specific threshold *D*_*t*_. We reformulated the problem as maximizing the harmonic mean of these two conditional probabilities. The harmonic mean is defined as:

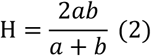

Given that both *a* and *b* are positive and their product *ab* = *constant*, the harmonic mean H reaches its maximum when (*a* + *b*) is minimized.

Considering the following equation:

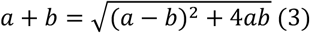

Under *ab* = *constant*, H is maximized when *a* − *b* = 0, i.e., when *a* = *b*. We assumed that intra-cluster and inter-cluster distance distributions are well separated, with substantially different means and minimal overlap.

Under this assumption, in an interval close to the point where H is maximized (*D*_*H*_), an increase in one conditional probability necessarily entails a decrease in the other. Let *f*_*a*_(*D*) and *f*_*b*_(*D*) denote the corresponding probability density functions. Then,

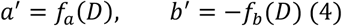

Because the overlap between the intra-cluster and inter-cluster distance distributions is assumed to be minimal, for an arbitrary sufficiently small positive constant 𝜀 (0 < 𝜀 ≪ 1), there exists an interval around *D*_*H*_ in which

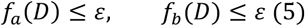

Meanwhile, since 0 ≤ *a* ≤ 1 and 0 ≤ *b* ≤ 1,

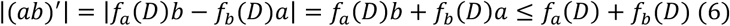

Around *D*_*H*_,

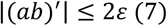

Thus, in the interval near *D*_*H*_, (*ab*)^′^ ≈ 0, so *ab* can be approximated as constant. Based on this estimation, the distance threshold can be inferred as a genomic distance value when the difference between the two conditional probabilities (*a* − *b*) is very close to zero. We defined the threshold as the distance at which (*a* − *b*) first becomes greater than zero, for all bins generated by dividing the 500bp to 100kb range into 500bp intervals.

The sparsity correction procedure consists of two steps:

First, we additionally merged two nearest enhancer candidates into the same cluster if they were included as enhancer candidates after gABC scoring, assigned to different enhancer clusters after distance threshold optimization, yet showed a very high cosine similarity, suggesting that they should reasonably belong to the same enhancer cluster. This approach is justified by the assumption that some enhancers are filtered during the gABC scoring process, and the inherent sparsity of scATAC-seq data can lead to this discrepancy.

Second, for peaks that were not included as enhancer candidates after gABC scoring but existed in the initial set of enhancer candidates before scoring, we reconsidered their inclusion if they (1) had a co-accessibility (correlation coefficient) greater than a specific threshold (default: 0.49) with at least one enhancer candidate and (2) showed very high accessibility. The default accessibility threshold was defined as the accessibility value of the top 0.5% of enhancer candidates after gABC scoring. Newly added enhancer candidates were assigned to the enhancer cluster of the linked enhancer candidate that exhibited the highest value of assignment score. Assignment score is defined as follows:

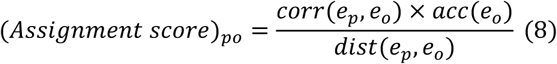

where *corr*(*e*_*p*_, *e*_*o*_) is the correlation coefficient between the newly added enhancer candidate *e*_*p*_ and the original enhancer candidate *e*_*o*_, *acc*(*e*_*o*_) is the accessibility of *e*_*o*_, and *dist*(*e*_*p*_, *e*_*o*_) is the genomic distance between *e*_*p*_ and *e*_*o*_.

### Pseudo-directed network construction

We assumed that the strength of the interaction between two nodes (two enhancer candidates) in the co-accessibility network is proportional to the co-accessibility (correlation coefficient), the inverse of the genomic distance, and the product of their accessibility values.

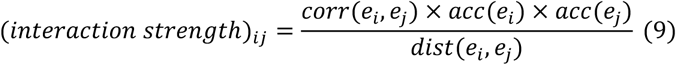

where *corr*(*e*_*i*_, *e*_*j*_) is the correlation coefficient between two different nodes *e*_*i*_ and *e*_*j*_, *acc*(*e*_*i*_) is the accessibility of *e*_*i*_, *acc*(*e*_*j*_) is the accessibility of *e*_*j*_, and *dist*(*e*_*i*_, *e*_*j*_) is the genomic distance between *e*_*i*_ and *e*_*j*_. Based on the co-accessibility network, we constructed a network with the above interaction strength as the network edge weight.

Inspired by the concept of electronegativity in chemistry, we assumed that the interaction strength assigned to each edge is asymmetrically distributed between two connected nodes in proportion to the total accessibility of the enhancer cluster to which each node belongs. To incorporate this weight into the calculation of node centrality (personalized PageRank ranking procedure), we transformed the constructed network from an undirected graph to a directed graph. Specifically, for each edge between node *e*_*i*_ and node *e*_*j*_, we redistributed the previously calculated interaction strength weights (as shown in Equation (9)) into two directed edges, one from *e*_*i*_ to *e*_*j*_ and the other from *e*_*j*_ to *e*_*i*_, based on the relative total accessibility of their corresponding enhancer clusters (Figure 1b). The network edge weight for an edge from *e*_*i*_ to *e*_*j*_ is as follows.

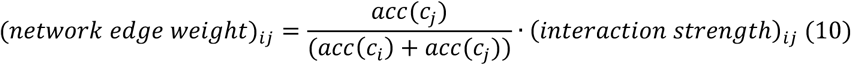

where *acc*(𝑐_*i*_) is the total summed accessibility of the enhancer cluster 𝑐_*i*_ corresponding to node *e*_*i*,_ and *acc*(𝑐_*j*_) is the total summed accessibility of the enhancer cluster 𝑐_*j*_ corresponding to *e*_*j*_. After assigning edge weights in the network, we further assigned node weights for the personalized PageRank calculation. The node weight is defined as follows:

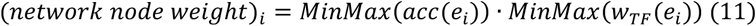

where 𝑀*i*𝑛𝑀*a*𝑥(*acc*(*e*_*i*_)) is the value set of 𝑀*i*𝑛𝑀*a*𝑥-scaled accessibility of node *e*_*i*_ across all nodes within the network, and 𝑀*i*𝑛𝑀*a*𝑥(𝑤_𝑇𝐹_ (*e*_*i*_)) is the value set of 𝑀*i*𝑛𝑀*a*𝑥-scaled TF weight of *e*_*i*_ across all nodes within the network. 𝑀*i*𝑛𝑀*a*𝑥-scaling was performed using the scales package (v1.3.0).

TF weight calculation for each node was performed as follows: First, we applied motifmatchr (v1.20.0) to identify the presence of CISBP TF motifs in each final enhancer candidate (enhancer constituent).

Subsequently, motif deviation scores (z-scores) computed per cell using ArchR (v1.0.2) were aggregated by calculating the median across all cells within the target cell population. The median z-score of each TF was defined as the motif enrichment weight (per cell population & per TF). TF motifs with motif enrichment weight below a pre-defined threshold (default: 0.675) were assigned a weight of 0. Finally, for each enhancer constituent, the motif enrichment weights for all present TF motifs were summed to obtain the node-specific TF weight.

### personalized PageRank ranking

For the constructed pseudo-directed network, we calculated personalized PageRank values using both node and edge weights using the igraph package (v1.5.1). Then, the PageRank values assigned to each node were aggregated per enhancer cluster, and the resulting sum was defined as the PageRank sum representing the relative importance of each enhancer cluster. For normalization, we scaled the maximum PageRank sum to 100 and ranked it in ascending order based on the scaled PageRank sum of the enhancer clusters.

In this context, the kneed package (v1.0.0) was adopted to determine the threshold of the scaled PageRank sum value corresponding to the elbow point of the ranked distribution. The region of the enhancer cluster where the scaled PageRank sum exceeds this threshold was defined as the Clustered Open Regulatory Elements (CORE). The initial sensitivity parameter was set to 2 to infer the elbow point. The threshold was accepted if the number of CORE regions identified using that elbow point did not exceed the maximum allowed number of CORE regions (default: 0.1*26000/1.31). Otherwise, the sensitivity parameter was repeatedly increased by 1, and the elbow points were recalculated until the number of CORE regions fell below the maximum limit. In some cases, constituents within the defined CORE may exhibit asymmetric chromatin accessibility, where regulatory activity is concentrated in only a subset of regions that cooperatively regulate similar gene expression programs. Consequently, this can result in overestimating the spatial and functional span of CORE. To address this problem, we excluded constituents with accessibility values below a specific threshold (default: 5) from the final CORE definition.

### Iterative proximal enhancer clusters filtering

We annotated CORE regions to their nearest genes using ChIPseeker (v1.34.1). The promoter accessibility of each annotated gene was defined as the sum of the normalized accessibility values of all peaks located within the promoter region (within 2kb from the TSS). Genes with promoter accessibility not exceeding 12.5 were considered to have inactive promoters. Subsequently, the genomic regions within a specific distance (default: 7.5kb upstream and downstream from the TSS of inactive promoters) were defined as inactive proximal enhancer regions. Using BEDTools (v2.31.0) subtract function, we subtracted inactive proximal enhancer regions from the raw CORE regions, resulting in a preliminary filtered set of CORE regions. The initially filtered CORE regions were then re-annotated to the nearest genes. If the initially filtered CORE regions were re-annotated as a gene with a TSS overlapping the inactive proximal enhancer region and was truncated in the previous first filtering step, these CORE regions were subsequently eliminated. After this additional filtering step, the remaining CORE regions were defined as the final output of the active option.

### scATAC-seq & scCUT&Tag-seq processing

Data retrieved from publicly available scATAC-seq datasets: 10k PBMC from 10X Genomics, BMMC (GSM4138889)^64^, CRC (GSE201349)^65^, and Fetal Lung (E-MTAB-11266; WSSS_F_LNG8782706 & WSSS_F_LNG8782707)^66^. ArchR (v1.0.2) was used for processing and analyzing scATAC-seq data. Low-quality cells were filtered based on the following quality measures: Cells with nCount_peaks > 3000 & nCount_peaks < 30000 & pct_reads_in_peaks > 15 & blacklist.ratio < 0.05 & nucleosome_signal < 4 & TSS.enrichment > 3 (PBMC); log10(nFrags) > 3.3 and TSSenrichment > 5 (BMMC); log10(nFrags) > 3.4 & TSSenrichment > 4 (CRC); log10(nFrags) > 3 & TSSenrichment > 4 (Fetal Lung) were retained. Doublets were detected and removed by using the ArchR filterDoublet function (filterRatio=1.0). The remaining cells were normalized and subjected to downstream analysis using the ArchR pipeline. Dimensionality reduction was performed using the Iterative Latent Semantic Indexing (IterativeLSI) method. Harmony method was applied to the reduced dimensions in order to remove unwanted non-biological variations. UMAP visualization was conducted based on the Harmony-corrected reduced dimensions. Imputed gene scores were calculated using MAGIC (v3.0.0).

With the exception of PBMC data, cell types were annotated based on MAGIC-imputed gene scores of canonical marker genes. For PBMC, to perform cell type annotation, the label transfer method was applied using Signac (v1.14.0) before ArchR processing. Normalized ATAC signals (by ReadsInTSS metric) were used to generate the genome browser track visualization result. Co-accessibility information was extracted by using the ArchR addCoAccessibility function with these parameters: corCutOff=0.2 and maxDist=500000.

For scCUT&Tag-seq analysis, we used publicly available data (GSE195725)^67^ for PBMC. The .rds files from GSE195725 were processed using Signac. Cell types were inferred based on clustering information and pre-annotated labels. Peaks were extracted for each cell type, and regions within 2kb of UCSC known genes’ TSS were excluded. The remaining peaks were stitched using a maximum distance threshold of 12.5kb. Finally, super-enhancers (SEs) were identified using the ROSE algorithm1.

### ChIP-seq processing

For validation of PBMC data, ChIP-seq datasets for H3K27ac, H3K4me1, and H3K27me3 were mainly collected from the ENCODE database^68^, except for CD16+ Monocytes. In the case of CD16+ Monocytes, H3K27ac ChIP-seq data were obtained from the FANTOM database (GSE40502)^69^. Raw FASTQ files were processed as follows: adapter trimming with Skewer (v0.2.2), hg19 genome alignment with Bowtie2 (v2.5.1), de-duplication with the PICARD (v2.27.4) MarkDuplicates command, and blacklist filtering with BEDTools (v2.31.0) intersect function. The blacklist-filtered BAM files were then used to construct Tag Directories with the HOMER (v4.11.1) makeTagDirectory command. For track visualization for CD14+ Monocytes, we used CD14+CD16- Monocytes data from GSE40502 and applied the same processing pipeline as described above. BedGraph files were generated with HOMER makeUCSCfiles function. Track visualization was performed using WashU Epigenome Browser^70^.

For validation of CRC data, two publicly available datasets were used: H3K27ac data from GSE156613^71^ and H3K27ac & H3K4me1 data from GSE136889^72^. To infer SE regions of CRC, 15 CRC samples were sub-sampled from each dataset (Supplementary Table 8). Additionally, 15 normal colon samples were chosen from GSE136889 for comparative analysis (Supplementary Table 8). Adapter trimming was performed using Skewer. Trimmed reads were aligned to the hg38 genome using Bowtie2. Duplicated reads were removed using the PICARD MarkDuplicates command. Blacklist regions were filtered using BEDTools intersect function. The blacklist-filtered BAM files were then used to construct Tag Directories with the HOMER makeTagDirectory command. For classification performance comparison between CORE and SE regions in CRC and normal colon samples, raw read count matrices were extracted using DiffBind (v3.8.4). Normalization was performed using DESeq2 (v1.38.3), with size factors calculated as library size divided by median library size across samples. For downstream analysis, rlog-transformed counts were used for Non-Negative Factorization (NMF; using NMF R package v0.28; random.seed=1-100), and normalized counts were used for XGBoost^73^ classification (after hyperparameter tuning via Grid Search with cv=5; random.seed=1-100 in iterative random holdout procedure with 0.75:0.25 train test split for test set validation).

SE calling was performed using HOMER findPeaks with the parameters -style super -L 0. Regions within 2kb of UCSC known genes’ TSS were excluded from SE identification using the -excludePeaks option.

### Bulk RNA-seq processing

To investigate cell type-specific transcriptional regulation of CORE-associated and SE-associated genes in PBMC, we utilized the Monaco PBMC RNA-seq dataset and the HPA PBMC RNA-seq dataset from the Human Protein Atlas (HPA; https://www.proteinatlas.org/)^74^. For gene expression values, we used normalized TPM values pre-calculated by the Human Protein Atlas. To quantify the contribution of CORE and SE to cell type-specific transcriptional regulation, we calculated the mean log2 fold changes (Specificity score) of normalized TPM values between CORE-associated genes vs non-CORE-associated genes or SE-associated genes vs TE-associated genes for each cell type based on the Monaco dataset.

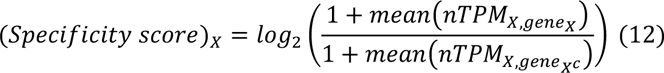

where *mean(nTPM_x,gene,x_)* is the average of normalized TPM measured in samples corresponding to cell type *X* for genes associated with region set derived from cell type *X*, and 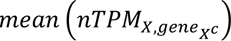 is the average of normalized TPM measured in samples corresponding to cell type *X* for genes associated with region set derived from other cell types *X^c^*.

To measure the overlap between differentially expressed genes (DEGs), CORE-associated genes, and SE-associated genes in CLL vs Normal B cells, we utilized the publicly available dataset (GSE66117)^75^. Raw read counts were subjected to RLE normalization using DESeq2 (v1.38.3). DEGs were identified as DESeq2 by applying a cut-off of |log2FoldChange| > 1 & q-value < 0.05. The log2 fold change values were shrunk using the apeglm package (v1.20.0). The degree of overlap between DEGs and CORE/SE-associated genes was quantified using the following formula:

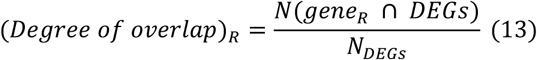

where 𝑁(𝑔*e*𝑛*e*_𝑅_ ∩ *D*𝐸𝐺𝑠) is the number of overlaps between DEGs and genes associated with region 𝑅, and 𝑁_*D*𝐸𝐺𝑠_ is the number of DEGs.

### Peak annotation & GO enrichment analysis

GREAT^76^ was used to identify CORE/non-CORE-associated and SE/TE-associated genes. For PBMC and Fetal Lung, peak-to-gene annotation was performed using the single nearest gene within 500kb of the TSS. For CRC, COREs were derived using the potential option, and each peak was assigned to two nearest genes within the same distance threshold to reduce missed peak-to-gene links in broad/multi-gene regulatory neighborhoods. For BMMC, gene linking was used only to examine recovery of representative hematopoietic lineage-defining transcription factors, or loci highlighted by the macrophage-monocyte differential H3K27ac analysis; a CORE was considered linked to a representative gene if the gene was either (i) the single nearest gene (within 500 kb) or (ii) the CORE midpoint fell within 8.25 kb of the gene’s TSS (after *Iterative proximal enhancer clusters filtering*, 7.5 kb promoter-proximal window used in the active option plus the midpoint of a stitched 3 x 501 bp peak unit), and these assignments were additionally validated by inspection of ArchR scATAC-seq tracks. GO enrichment analysis for CD14+ and CD16+ Monocytes of PBMC was also performed using GREAT.

### Chromatin interaction enrichment analysis

For H3K27ac HiChIP data analysis, FitHiChIP 5k loop information for GM12878, THP-1, and Th17 was obtained from HiChIPdb^77^. Using loop information, we constructed the chromatin interaction network with the igraph package (v1.5.1) and calculated weighted degree (strength) for each node. Normalized count values were used as network edge weights.

For Hi-C data analysis, loop information (BEDPE files containing HiCCUPS outputs) for B cells (ENCSR847RHU) and CD14+ Monocytes (ENCSR236EYO) was obtained from the ENCODE database^68^. Loops with FDR > 0.01 were removed (observed/expected ratio-based FDR). After that, only the loops in which the two interaction regions overlapped with the H3K4me1 peaks of the corresponding cell type (CD19+ B cells: ENCSR214VUB, CD14+ Monocytes: ENCSR400VWA) were retained. We then constructed the enhancer-enhancer chromatin interaction network using igraph and calculated weighted degree (strength) for each node. Network edge weights were defined as observed-minus-expected counts to quantify contact signals above the local expected background. This metric was used to preserve differences in interaction magnitude, as constituent-level interaction strength derived from ratio-based measures can be attenuated in interaction-dense neighborhoods with elevated local expected counts. As the Hi-C data were provided in hg38, CORE and SE coordinates originally defined in hg19 were converted to hg38 using UCSC LiftOver^78^.

To characterize the epigenetic states of enhancer constituents within CORE and SE, we categorized H3K4me1-marked enhancer peaks into three state categories: active, poised, and primed enhancers. The detailed procedure is as follows: Among the ChIP-seq datasets described in the *ChIP-seq processing* section above, we used the H3K27ac for CD19+ B cells (ENCSR191ZQT) and CD14+ Monocytes (ENCSR012PII), and the H3K27me3 for CD19+ B cells (ENCSR404MOX) and CD14+ Monocytes (ENCSR080XUB) from the ENCODE database^68^. Read count matrices were generated for H3K27ac and H3K27me3 at pre-defined H3K4me1 peaks using the DiffBind (v3.8.4). The data was normalized with CPM (Counts per Million). H3K4me1 peaks were classified into three state categories based on the log2 fold change between H3K27ac and H3K27me3 signals: peaks with a value greater than 1 were classified as active, those with a value less than -1 as poised, and the remaining peaks as primed.

### GWAS enrichment analysis

For GWAS enrichment analysis, in the present study, we collected GWAS variants associated with immune-related diseases and non-immune-related traits through two approaches. First, we utilized the fine-mapped GWAS variants (95% credible set from FINEMAP metric) curated from several GWAS cohorts via CausalDB^79^. Second, to facilitate direct comparison with SE, we used the Young Lab-curated version of GWAS variants (Hnisz *et al.*) that first reported the correlation between SE and disease-related GWAS enrichment^2^. To measure GWAS enrichment for CORE and SE, we defined the enrichment score as follows:

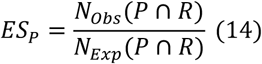

where 𝑁_𝑂*b*𝑠_(*P* ∩ 𝑅) is the observed number of GWAS variants included in the GWAS phenotypic trait *P* within region set 𝑅, and 𝑁_𝐸𝑥*p*_(*P* ∩ 𝑅) is the expected number of GWAS variants included in the GWAS phenotypic trait *P* within a randomly permuted region set derived from region set 𝑅 with 1000 iterations.

To measure cell type specificity for fine-mapped CAD-risk loci enrichment, the specificity index was defined as follows.

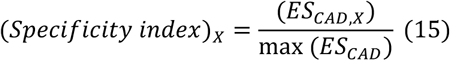

where 𝐸𝑆_𝐶𝐴*D*,𝑋_ is the GWAS enrichment score for fine-mapped CAD risk loci in cell type 𝑋, and max (𝐸𝑆_𝐶𝐴*D*_) is the maximum value of the GWAS enrichment scores from all cell types, including 𝑋.

### TF motif enrichment analysis

To perform both known motif enrichment and *de novo* motif enrichment analyses, the HOMER (v4.11.1) findMotifsGenome.pl function was used with the -size given & -mask parameters. For PBMC data, only the known motif enrichment analysis results were adopted. Calculation of the motif enrichment score was performed using the modified MinMax scaling method as follows.

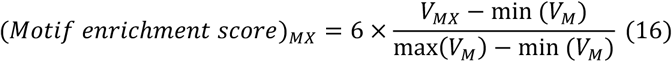

where 𝑉_𝑀𝑋_ is the statistical significance (-log10(P-value)) for TF motif 𝑀 within CORE regions from any cell type 𝑋, max(𝑉_𝑀_) is the maximum 𝑉_𝑀𝑋_ across all cell types, and min (𝑉_𝑀_) is the minimum 𝑉_𝑀𝑋_ across all cell types. Modified 𝑀*i*𝑛𝑀*a*𝑥-scaling was performed using the scales package (v1.3.0).

Only significant motifs with p-value < 1e-04 & q-value < 0.05 were included in the known motif enrichment analysis. For CRC, the results from the *de novo* motif enrichment analysis were used. Significant motifs for *de novo* motif enrichment analysis were defined as those not marked by possible false positives.

### Similarity network construction for bone marrow and peripheral blood hematopoietic cells

To incorporate the CORE regions derived from each BM cell type, we first employed the BEDTools (v2.31.0) sort and merge function. We used the BEDTools intersect function to determine whether the merged regions overlapped with the CORE regions of each cell type. Overlaps were binarized as 1 (overlap) or 0 (no overlap), resulting in a binary region x cell type matrix format. Based on the binary matrix, we calculated the pairwise cosine similarity between cell types using the coop package (v0.6-3). We then constructed the network with the igraph package (v1.5.1) using the cosine similarity matrix. Edges with cosine similarity < 0.55 were removed to refine the network. Network visualization was performed using the ggraph package (v2.2.1). For network construction on BM-to-PB maturation ordering, the same procedure was applied to PB cell types. Pairwise cosine similarities were calculated between BM and PB cell types. Then, edges with cosine similarity < 0.55 were removed to refine the BM-to-PB bipartite network.

### Single-cell Multiome velocity analysis

We analyzed the single cell multiome dataset for CD34+ hematopoietic cells provided in the original MultiVelo paper and tutorial^80^. Cell types were inferred based on pre-annotated labels. For Multiome velocity estimation, we used the following parameters: n_pc=10, n_neighbors=20, weight_c=0.75.

### Enrichment analysis of H3K27ac differential peaks between macrophages and monocytes

Raw FASTQ files for H3K27ac profiles of macrophages and monocytes were retrieved from publicly available datasets (GSE31621)^81^, together with a macrophage/monocyte H3K4me1 peak set used for downstream quantification. Then, using the same pipeline as above *ChIP-seq processing* for PBMC, reads were aligned to the hg19 reference genome. Both blacklist-filtered BAM files and BedGraph files were generated. Using DiffBind (v3.8.4), raw read count matrices for H3K27ac profiles were extracted using H3K4me1 peaks derived from macrophages and monocytes as the peak set. CPM (Counts per million) normalization was applied to the raw read count matrices. Differential peaks between macrophages and monocytes were obtained using |log2FoldChange| > 3 as a cut-off value. The Cochran-Mantel-Haenszel (CMH) test was used for enrichment analysis. The detailed procedure is as follows: To compare CORE and SE enrichment, we used the regions where TEs do not overlap any COREs or SEs as a control peak set. Peak widths for the three types of peak set (CORE, SE, and TE) were binned to quantile ranges. With the generated bin as strata, the CMH test was applied to contingency tables summarizing the overlap between differential peaks (macrophage-elevated, monocyte-elevated) and each peak set comparison (COREs vs TEs, SEs vs TEs), yielding common odds ratios and p-values. Differential peaks within 2kb of UCSC known genes’ TSS were excluded for enrichment analysis. The Breslow-Day (BD) test was used as a homogeneity test to determine the reliability of the calculated common odds ratios. For all comparisons, the common odds ratio was determined to be confident by satisfying p-value > 0.05 derived from the BD test. Track visualization for representative genes (*MEF2A*, *BHLHE41*, and *DCSTAMP*) was performed using WashU Epigenome Browser^70^. Differential peaks near BHLHE41 were visualized but excluded from the enrichment analysis because they were located within 2kb of TSS.

### TCGA survival analysis

Survival analysis for CORE-associated genes was performed using the Colon Adenocarcinoma (COAD) cancer subtype data from the TCGA database. Survival curves were produced using the survival package (v3.4-0). Statistical significance P values were computed using a log-rank test.

### Enrichment analysis for CRC master TF candidates

Master TF candidates for CRC were collected from previous reports^59,82–84^. The enrichment score was defined as the proportion of CRC master TF candidates that overlap with CORE- or SE-associated genes using the following formula:

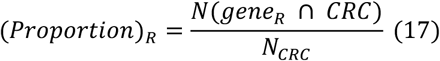

where 𝑁(𝑔*e*𝑛*e*_𝑅_ ∩ 𝐶𝑅𝐶) is the number of overlaps between CRC master TF candidates and genes associated with region 𝑅, and 𝑁_𝐶𝑅𝐶_ is the number of CRC master TF candidates.

### Network-based analysis and Metastasis-related genes

Based on the co-accessibility network information of epithelial cells in the CRC scATAC-seq dataset^65^, an undirected graph was constructed using the igraph package (v1.5.1), with correlation coefficients assigned as network edge weights. To focus on enhancer-associated elements, nodes were restricted to peaks overlapping CRC H3K4me1 peaks. Peaks within 2kb of UCSC known genes’ TSS were excluded. We then calculated weighted degree (strength) and local clustering coefficient (transitivity) for each node. Metastasis-related TFs were obtained from the COAD-related single-cell RNA-seq-derived TF regulon provided by metsDB^85^. Metastasis-related genes were identified from the TCGA GDC using cBioPortal^86^. The detailed procedures for extracting metastasis-related genes are as follows: Raw read counts were subjected to RLE normalization using DESeq2 (v1.38.3). Differential expression analysis was then performed between M1 (metastasis) and M0 (no metastasis). Metastasis-related genes were identified by applying a cut-off of log2FoldChange > 0.5 & q-value < 0.01. The abundance of metastasis-related TFs and metastasis-related genes was quantified using the following formula:

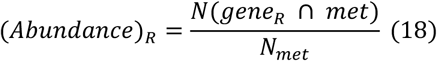

where 𝑁(𝑔*e*𝑛*e*_𝑅_ ∩ 𝑚*et*) is the number of overlaps between metastasis-related TFs or metastasis-related genes and genes associated with region 𝑅, and 𝑁_𝑚*et*_ is the number of metastasis-related TFs or metastasis-related genes.

Additionally, to measure the enrichment of metastasis-related TFs and metastasis-related genes within CORE, relative fold changes for the number of peak-annotated genes that overlap with metastasis-related TFs or metastasis-related genes were calculated as follows.

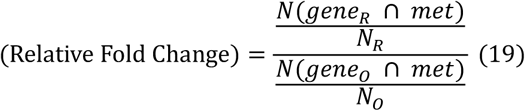

where 𝑔*e*𝑛*e*_𝑅_ is the gene set associated with peaks within COREs, 𝑔*e*𝑛*e*_𝑂_ is the gene set associated with peaks without COREs, 𝑁(𝑔*e*𝑛*e*_𝑅_ ∩ 𝑚*et*) is the number of overlaps between metastasis-related TFs or metastasis-related genes and genes within 𝑔*e*𝑛*e*_𝑅_, 𝑁(𝑔*e*𝑛*e*_𝑂_ ∩ 𝑚*et*) is the number of overlaps between metastasis-related TFs or metastasis-related genes and genes within 𝑔*e*𝑛*e*_𝑂_, 𝑁_𝑙_ is the number of genes within 𝑔*e*𝑛*e*_𝑂_, 𝑁_𝑅_ is the number of genes within 𝑔*e*𝑛*e*_𝑅_, and 𝑁_𝑂_ is the number of genes within 𝑔*e*𝑛*e*_𝑂_.

Cancer stem cell (CSC)-associated genes were obtained from the clinical biomarker list for Colon cancer in BCSCdb^87^. The abundance of CSC-associated genes was calculated using the same procedure as described for Equation (18).

### *In silico* enhancer knockout analysis

To perform *in silico* enhancer knockout analysis, we used AlphaGenome (v0.0.1)^88^ with the Left colon ontology (UBERON0008971). We first selected candidate enhancer peaks from the CRC H3K27ac profiles^72^ by extracting typical enhancer peaks that overlapped the USP7-proximal CORE. To minimize unwanted effects related to transcript structure (e.g., splicing), we restricted the analysis to peaks annotated as distal intergenic. The DNA sequence corresponding to the selected peak was retrieved using the getSeq function in the BSgenome package (v1.66.3) and used as the reference variant. Enhancer knockout was modeled by setting the alternative variant to a deletion (REF = upstream base + deleted sequence; ALT = upstream base), and the predict_variant function was used to predict the knockout-associated changes in the Left colon RNA-seq profile. The prediction interval was defined as a 1Mb window centered on the USP7 locus (SEQUENCE_LENGTH_1MB). To measure variant effects, variant scores were computed for each gene under the Left colon ontology using the tidy_scores function. As independent chromatin contact evidence, we incorporated publicly available H3K27ac HiChIP data from the HT29 cell line (GEO: GSM7117686)^89^. Intra-chromosomal loop coordinates provided in hg19 were converted to hg38 using UCSC LiftOver^78^. The lifted-over loops were visualized together with CRC & Normal H3K27ac ChIP-seq^72^ signal tracks using the WashU Epigenome Browser^70^.

### Identification of eNet hub genes

eNet hub enhancers were defined as enhancers classified by the eNet^60^ algorithm as network hubs or module hubs. eNet hub genes were defined as genes assigned to at least one eNet hub enhancer, or genes corresponding to enhancer clusters labeled as Complex or Modular. For Fetal Lung, two sources were used: (1) eNet hub genes curated in eNetDB (Human Fetal Lung) and (2) eNet hub genes obtained by applying eNet2.0 to the Fetal Lung scATAC-seq dataset (E-MTAB-11266) using default parameters. For CRC, eNet hub genes and hub enhancers curated in eNetDB (CRC Disease) were used.

### Statistics and reproducibility

No statistical method was adopted to predetermine sample size. The experiments in the present study were not randomized, as all datasets are publicly available from observational studies. In case of the XGBoost classification, the training set and test set were randomly selected, and this analysis was repeated 100 times. The investigators were not blinded to allocation during experiments and outcome assessment. For hypothesis testing in a single group, Statistical significance P values were computed using a two-sided Wilcoxon signed-rank, Wilcoxon rank sum test. For multiple group comparison, P values were calculated using Dunn’s test as a post-hoc test after the Kruskal-Wallis test. Multiple hypothesis testing correction was performed by the Holm correction method.

## Data availability

The publicly available datasets were downloaded via Gene Expression Omnibus; scATAC-seq (GSM4138889, GSE201349), scCUT&Tag-Pro (GSE195725), ChIP-seq (GSE40502, GSE156613, GSE136889, GSE31621), RNA-seq (GSE66117), and HiChIP (GSM7117686). scATAC-seq dataset for 10X PBMC 10k was retrieved from a web repository (https://cf.10xgenomics.com/samples/cell-atac/1.0.1/atac_v1_pbmc_10k/atac_v1_pbmc_10k_fragments.tsv.gz). scATAC-seq dataset for Fetal Lung was acquired from the Human Lung Cell Atlas (E-MTAB-11266). Super-enhancer and typical enhancers for PBMC were downloaded from SEdb^19^. The Monaco PBMC RNA-seq dataset and the HPA PBMC RNA-seq dataset were obtained from the Human Protein Atlas (HPA; https://www.proteinatlas.org/). Hi-C loop information and ChIP-seq for H3K27ac, H3K27me3, and H3K4me1 (CD4+ T cells, CD8+ T cells, NK cells, B cells, and CD14+ Monocytes) were collected from the ENCODE database^68^. HiChIP for H3K27ac (GM12878, THP-1, and Th17) were obtained from HiChIPdb^77^. This work used fine-mapped and curated GWAS variants for immune-related diseases and non-immune traits. Fine-mapped GWAS variants used in this paper were downloaded from CausalDB^79^. Young Lab-curated version of GWAS variants used in this paper are available through the original publication with PMID: 24119843. Single cell multiome dataset for CD34+ hematopoietic cells are available in the MultiVelo tutorial (https://github.com/welch-lab/MultiVelo). Metastasis-related TFs and metastasis-related genes were obtained from metsDB^85^ and TCGA GDC.

## Code availability

enCORE is available on GitHub at https://github.com/R-Krait/enCORE. Code for the PBMC 3k demo dataset can be found at https://figshare.com/articles/dataset/Demo_PBMC_3k_enCORE_/31577779.

## Contributions

S.P. and S.H.P. conceptualized the study. S.P. contributed to the design and implementation of the method and performed all computational analyses. S.P., S.M., W.L., and S.H.P. contributed to the interpretation of the results. S.P. and S.H.P. wrote the original draft, and all authors contributed to manuscript review and editing. All authors read and approved the final version of the manuscript.

---

**Extended Data Fig.1:**
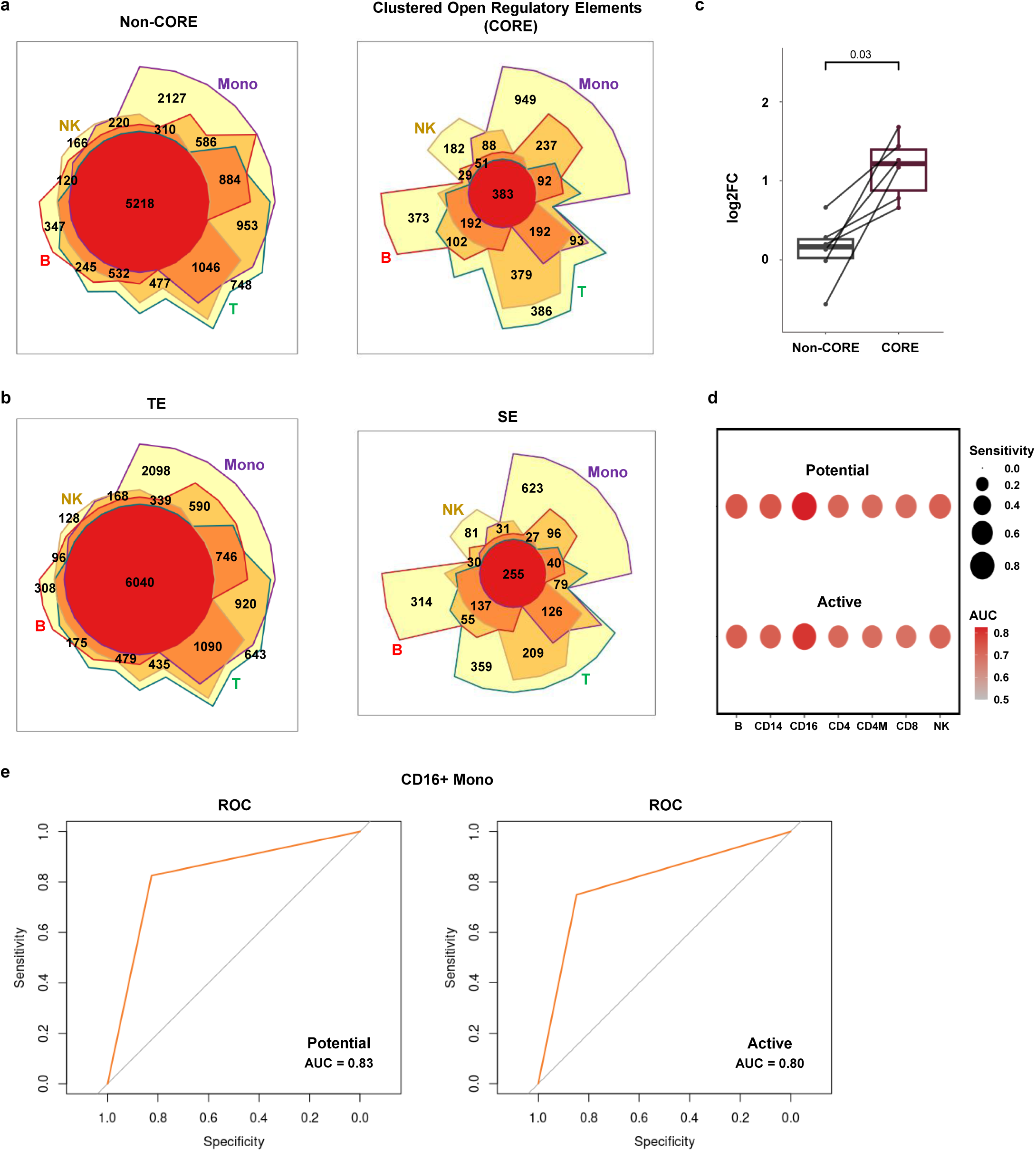
Cell type-specific transcriptional regulation captured by potential-option COREs a-b, Chow-Ruskey diagram of **a,** non-CORE-associated genes and CORE-associated genes; **b,** TE-associated genes and SE-associated genes in PBMC. B cells (red border), Monocytes (purple border), T cells (green border), NK cells (brown border). Monocytes represent the union of region-annotated genes derived from CD14+ and CD16+ Monocytes, while T cells represent the combined set of region-annotated genes derived from CD4+ and CD8+ T cells. The color of the borders around each intersection corresponds to the cell types whose genes overlap. The area of each intersection is proportional to the number of genes within the intersection. **c,** Log2-transformed fold changes of normalized TPM of genes in each cell type. Normalized TPM values were retrieved from the PBMC Monaco dataset in the Human Protein Atlas (HPA). Comparison between non-CORE-associated genes and CORE-associated genes using the potential option. P-value was calculated by a two-sided Wilcoxon signed-rank test. **d,** Dot plot showing the classification performance of CORE in distinguishing SE and TE. Dots are colored by Area Under Curve (AUC), and the dot sizes indicate the specificity. SEs derived from B: B cells, CD14: CD14+ Monocytes, CD16: CD16+ Monocytes, CD4: CD4+ T cells, CD4M: CD4+ Memory T cells, CD8: CD8+ T cells, NK: NK cells. **e,** Representative receiver operating characteristic (ROC) curves from CD16+ Monocytes. Left: COREs from the potential option, Right: COREs from the active option. The AUC values are indicated in the lower right corner of each plot.

**Extended Data Fig.2:**
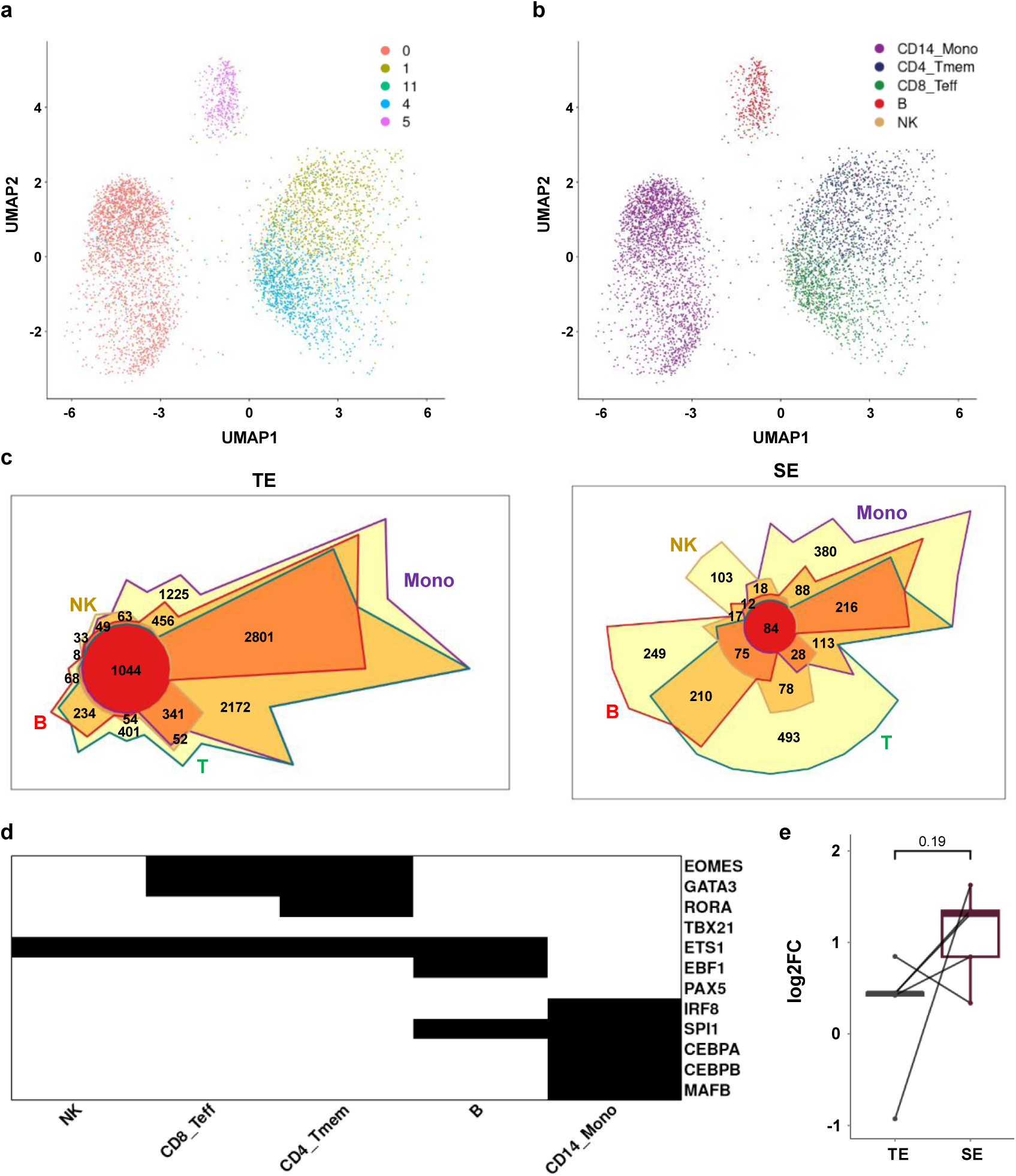
Benchmarking enCORE against SEs derived from PBMC H3K27ac scCUT&Tag-seq data a, UMAP embedding of the PBMC scCUT&Tag-seq dataset, colored and annotated by clusters. **b,** UMAP embedding of the PBMC scCUT&Tag-seq dataset, colored and annotated by cell types. CD14_Mono: CD14+ Monocytes, CD4_Tmem: CD4+ Memory T cells, CD8_Teff: CD8+ Effector T cells, B: B cells, NK: NK cells. **c,** Chow-Ruskey diagram of TE-associated genes and SE-associated genes. B cells (red border), Monocytes (purple border), T cells (green border), NK cells (brown border). T cells represent the combined set of SE-associated genes derived from CD4+ and CD8+ T cells. The color of the borders around each intersection corresponds to the cell types whose genes overlap. The area of each intersection is proportional to the number of genes within the intersection. **d,** Heatmap for cell type-specific master regulators. Black: gene present in the SE-associated genes of the given cell type. White: absent. **e,** Log2-transformed fold changes of normalized TPM of genes in each cell type. Normalized TPM values were retrieved from the PBMC Monaco dataset in the Human Protein Atlas (HPA). Comparison between TE-associated genes and SE-associated genes. P-value was calculated by a two-sided Wilcoxon signed-rank test.

**Extended Data Fig.3:**
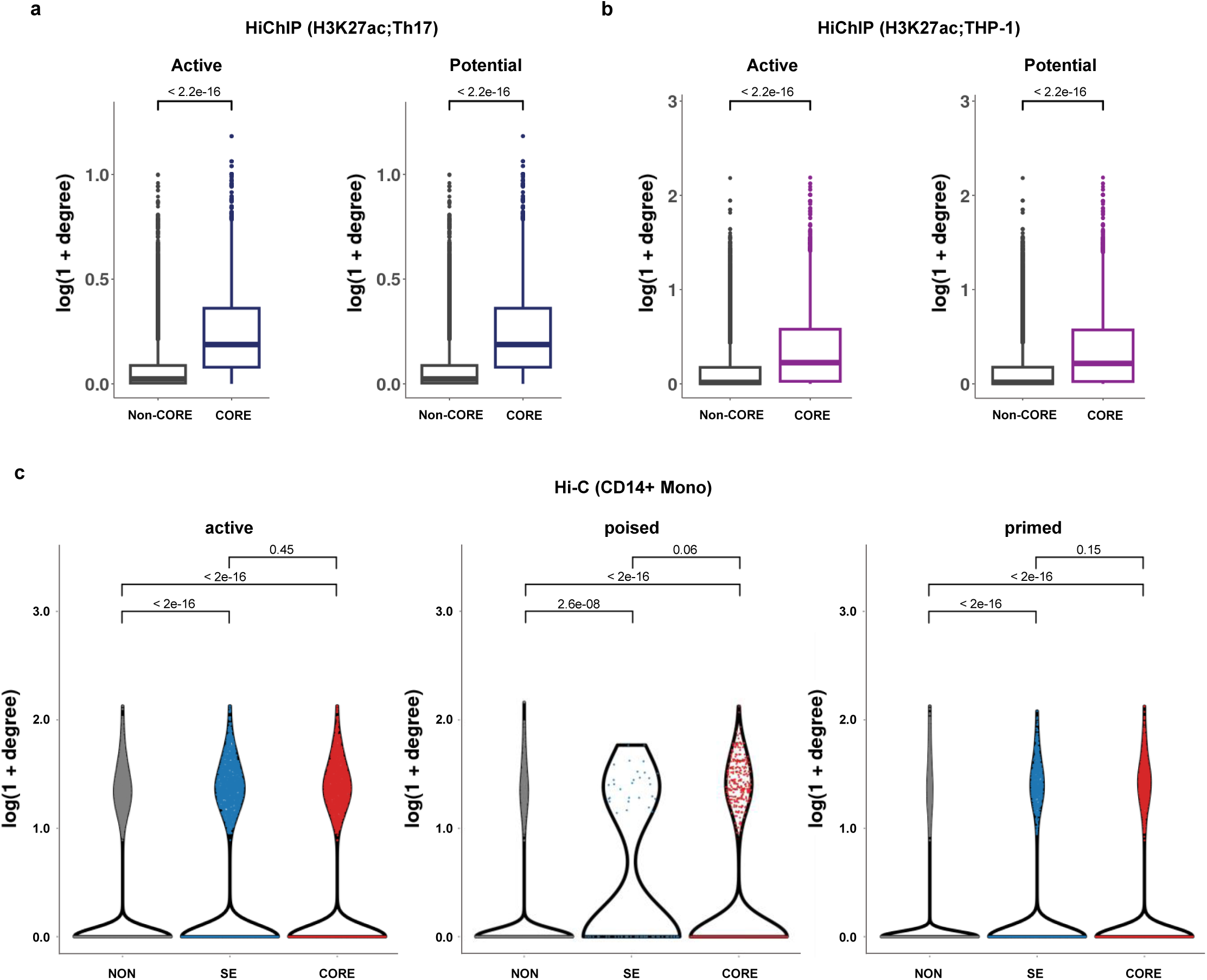
Enrichment of chromatin interactions within COREs in diverse cell types a, Log10-transformed weighted degrees of constituents within COREs and non-COREs. Data was retrieved from the H3K27ac HiChIP dataset for Th17 cells. **b,** Log10-transformed weighted degrees of constituents within COREs and non-COREs. Data was retrieved from the H3K27ac HiChIP dataset for THP-1 cells. Left: COREs from the active option, Right: COREs from the potential option. **c,** Log10-transformed weighted degrees of H3K4me1 peaks in each enhancer category from CD14+ Monocytes. Left: active enhancers, Middle: poised enhancers, Right: primed enhancers. Peak group marked by NON represents H3K4me1 peaks that do not fall into either COREs or SEs. Data was retrieved from the Hi-C dataset for CD14+ Monocytes. Only Hi-C interactions overlapping with H3K4me1 peaks are considered. COREs from the potential option. P-values were calculated to measure statistical significance (**a,b**, two-sided Wilcoxon rank sum test; **c**, Kruskal-Wallis test with Dunn’s test. P-values were adjusted by the Holm correction method).

**Extended Data Fig.4:**
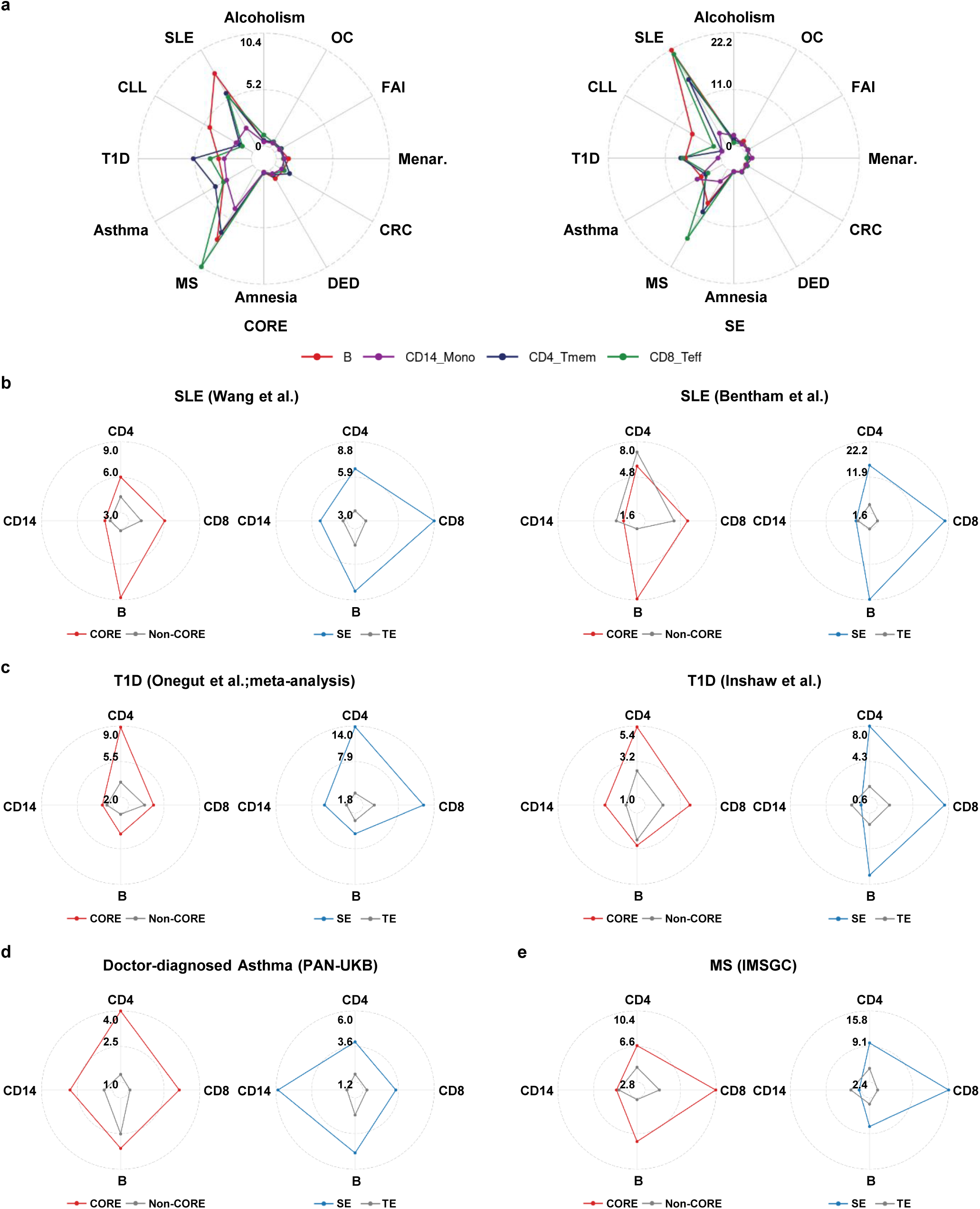
Robust enrichment of fine-mapped immune-related GWAS variants within COREs across multiple cohorts a, Enrichment analysis of fine-mapped GWAS variants for immune-related diseases or non-immune-related traits in COREs (left) and SEs (right). SLE: Systemic Lupus Erythematosus, CLL: Chronic Lymphocytic Leukemia, T1D: Type 1 Diabetes, Asthma: Asthma, MS: Multiple Sclerosis, Amnesia: Amnesia, DED: Dry Eye Disease, CRC: Colorectal Cancer, Menar.: Menarche, FAI: Fasting Insulin, OC: Ovarian Cancer, Alcoholism: Alcoholism. **b-d,** Enrichment analysis of fine-mapped GWAS variants for **b,** SLE; **c,** T1D; **d,** Asthma; **e,** MS in COREs and SEs. Lines are colored by region categories: COREs (red color), SEs (blue color), and TEs (grey color). B: B cells, CD14: CD14+ Monocytes, CD4: CD4+ Memory T cells, CD8: CD8+ Effector T cells.

**Extended Data Fig.5:**
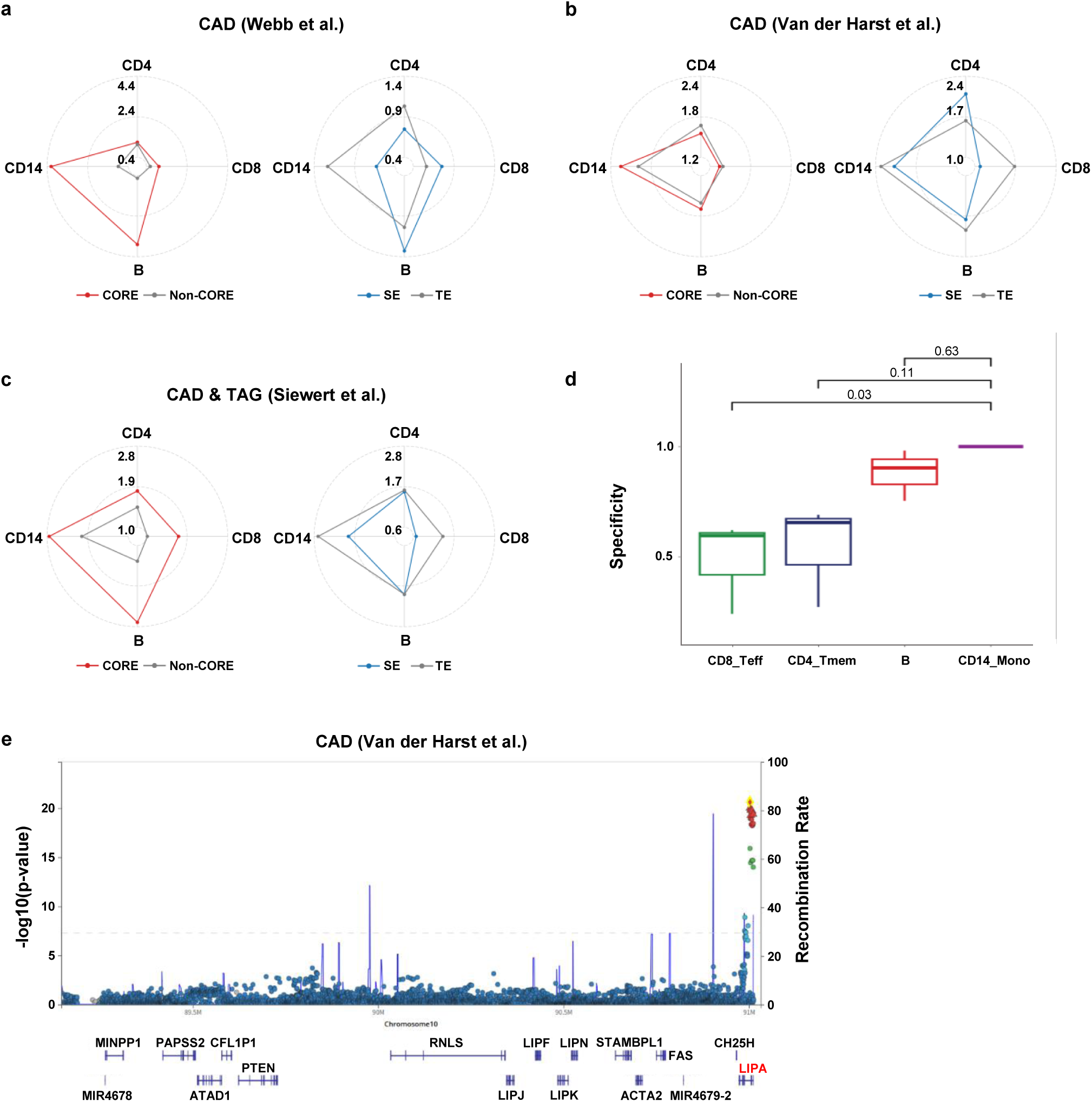
Enrichment of fine-mapped CAD risk loci within COREs a, Enrichment analysis of fine-mapped coronary artery disease (CAD) risk loci from Webb et al. GWAS cohort in COREs and SEs. **b,** Enrichment analysis of fine-mapped CAD risk loci from Van der Harst et al. GWAS cohort in COREs and SEs. **c,** Enrichment analysis of fine-mapped multi-variate CAD and TAG risk loci from Siewert et al. GWAS cohort in COREs and SEs. Lines are colored by region categories: COREs (red color), SEs (blue color), and TEs (grey color). **d,** Cell type specificity of fine-mapped CAD risk loci enrichment. Borderlines are colored by cell types. **e,** Fine-mapped CAD risk loci near the *LIPA* gene. The *LIPA* gene is highlighted in red. Cell types were indicated as abbreviations (**a-d**, B: B cells, CD14: CD14+ Monocytes, CD4: CD4+ Memory T cells, CD8: CD8+ Effector T cells).

**Extended Data Fig.6:**
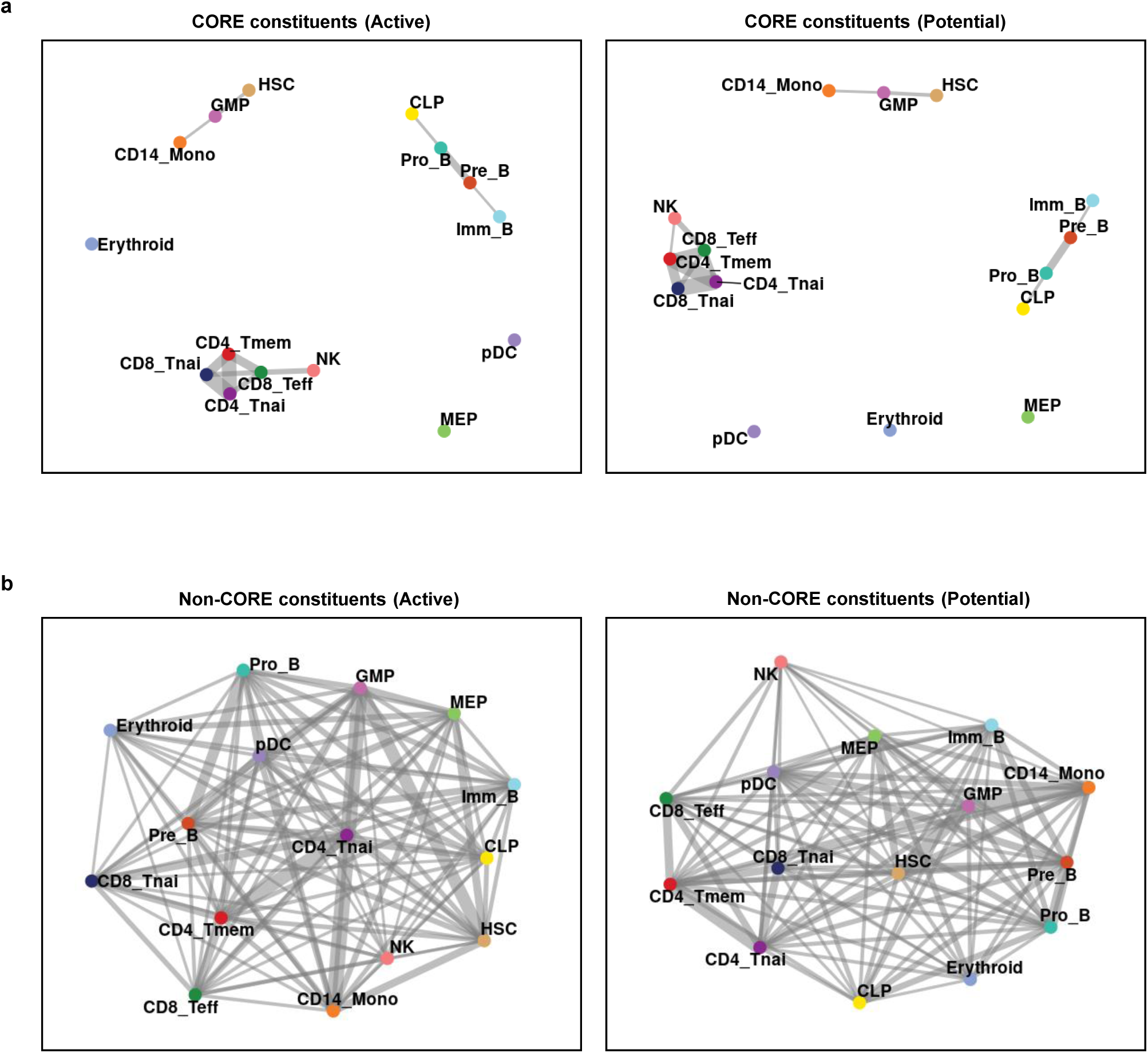
CORE constituents delineate lineage-specific epigenetic states a, Graph-based representation of the similarity among CORE constituents identified across different cell types in BMMC. Left: CORE constituents from the active option. Right: CORE constituents from the potential option. **b,** Graph-based representation of the similarity among non-CORE constituents identified across different cell types in BMMC. Left: non-CORE constituents from the active option. Right: non-CORE constituents from the potential option. Nodes are colored and annotated by cell types. Edges are shown only for node pairs with cosine similarity greater than 0.55. Cell types were indicated as abbreviations (**a,b**, HSC: Hematopoietic Stem Cells, GMP: Granulocyte-Myeloid Progenitors, CLP: Common Lymphoid Progenitors, MEP: Megakaryocyte-Erythroid Progenitors, CD14_Mono: CD14+ Monocytes, pDC: plasmacytoid Dendritic cells, Pro_B: Pro-B cells, Pre_B: Pre-B cells, Imm_B: Immature B cells, Erythroid: Erythroid cells, NK: NK cells, CD8_Teff: CD8+ Effector T cells, CD8_Tnai: CD8+ naïve T cells, CD4_Tmem: CD4+ Memory T cells, CD4_Tnai: CD4+ naïve T cells).

**Extended Data Fig.7:**
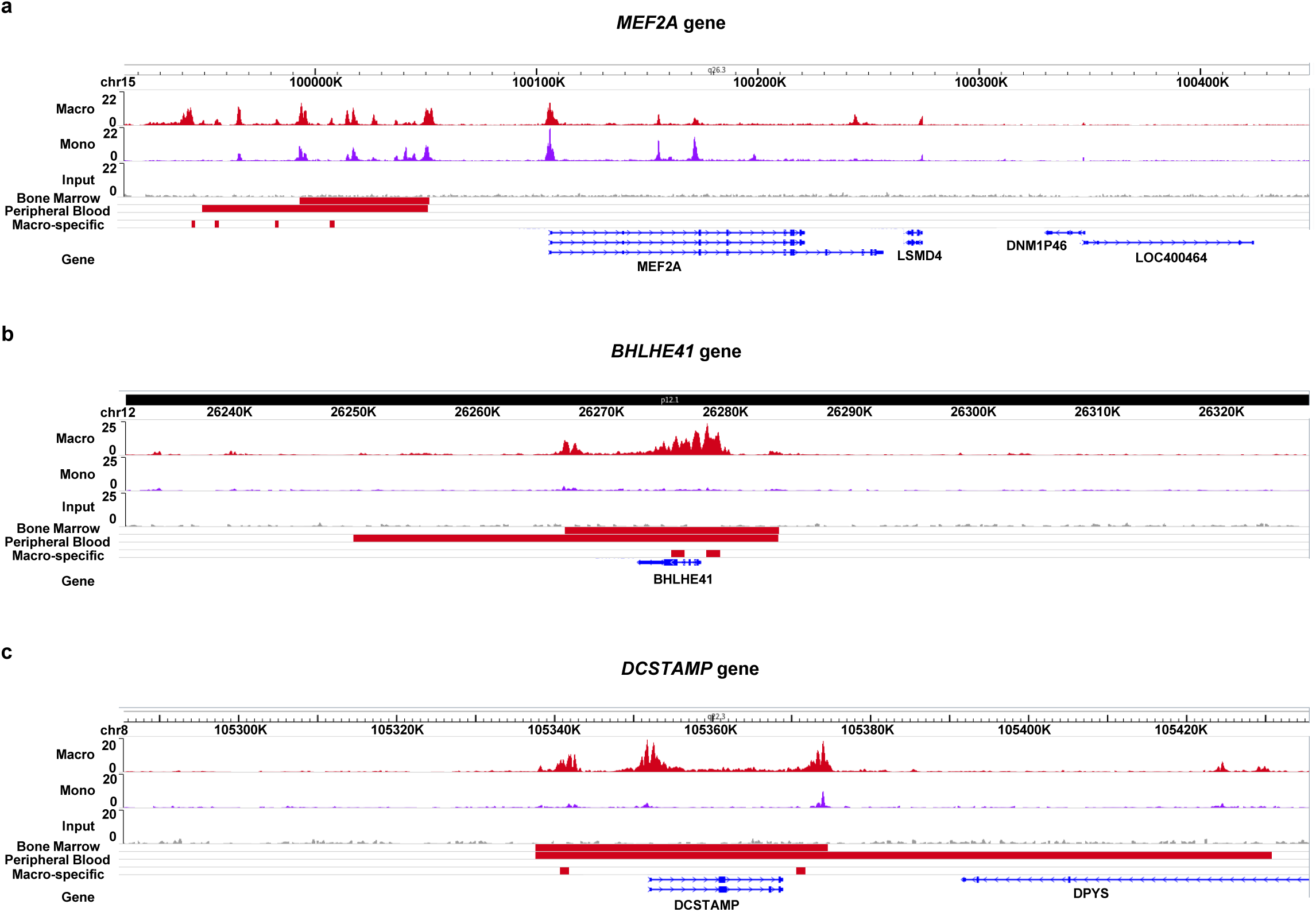
Track visualization of MEF2A, BHLHE41, and DCSTAMP in macrophages a, Track visualization at the *MEF2A* locus with normalized H3K27ac ChIP-seq signals in macrophages and monocytes. **b,** Track visualization at the *BHLHE41* locus with normalized H3K27ac ChIP-seq signals in macrophages and monocytes. **c,** Track visualization at the *DCSTAMP* locus with normalized H3K27ac ChIP-seq signals in macrophages and monocytes. Bone marrow: COREs from BM CD14+ Monocytes using the potential option. Peripheral blood: COREs from PB CD14+ Monocytes using the potential option. Macro-specific: macrophage-elevated H3K27ac peaks without TSS exclusion. Colored and annotated by cell types: Macro (Macrophages; red color), Mono (Monocytes; purple color).

**Extended Data Fig.8:**
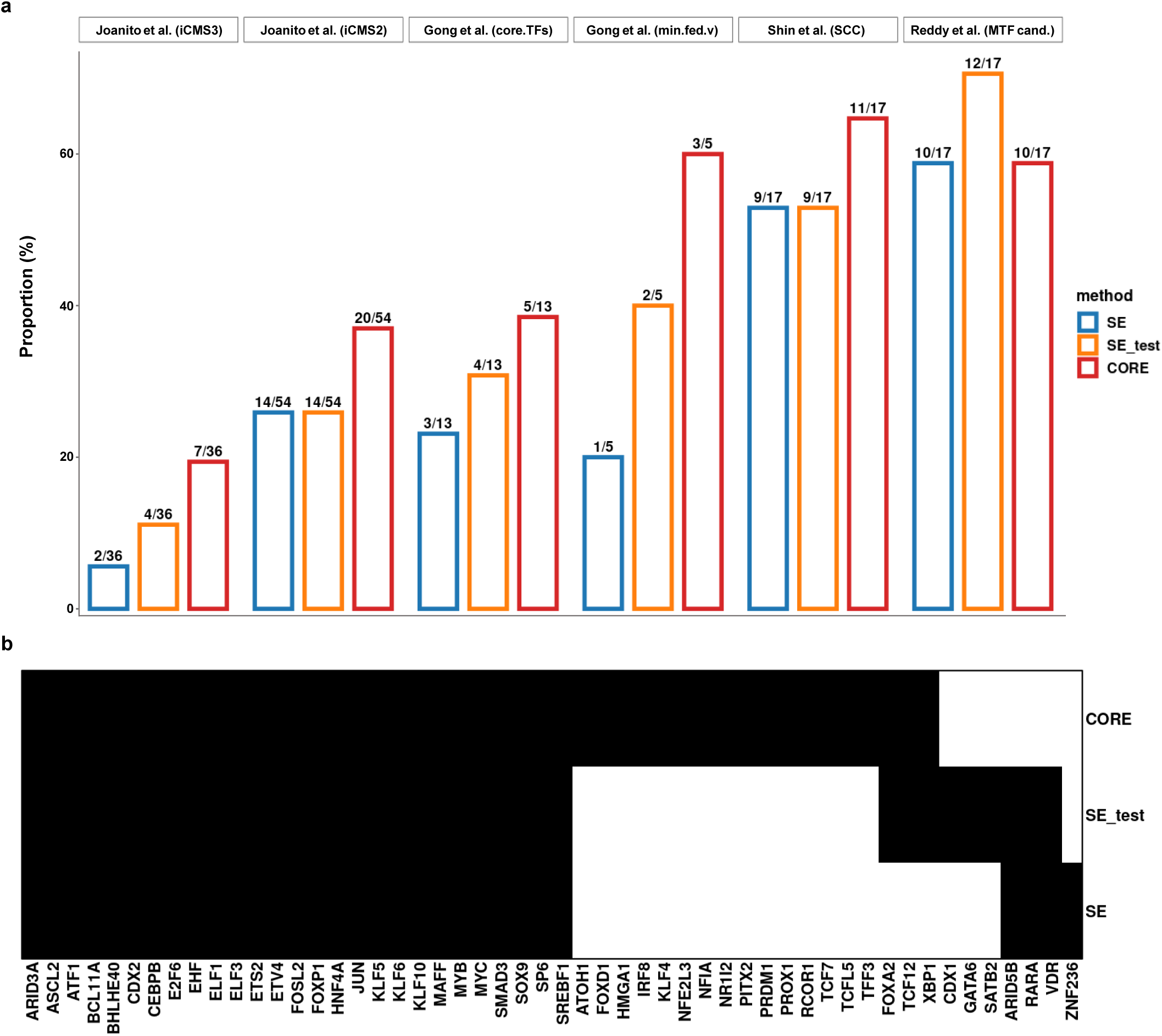
CRC master transcription factors are enriched among CORE-associated genes a, Proportion of CRC master TF candidates that overlap genes associated with each region category, relative to the total number of CRC master TF candidates in each study. Region category: COREs (CORE; red color), SEs from GSE136889 (SE_test; orange color), and SEs from GSE156613 (SE; blue color). **b,** Heatmap of CRC master TF candidates. Black: gene present in the region category-associated genes of the given cell type. White: absent.

**Extended Data Fig.9:**
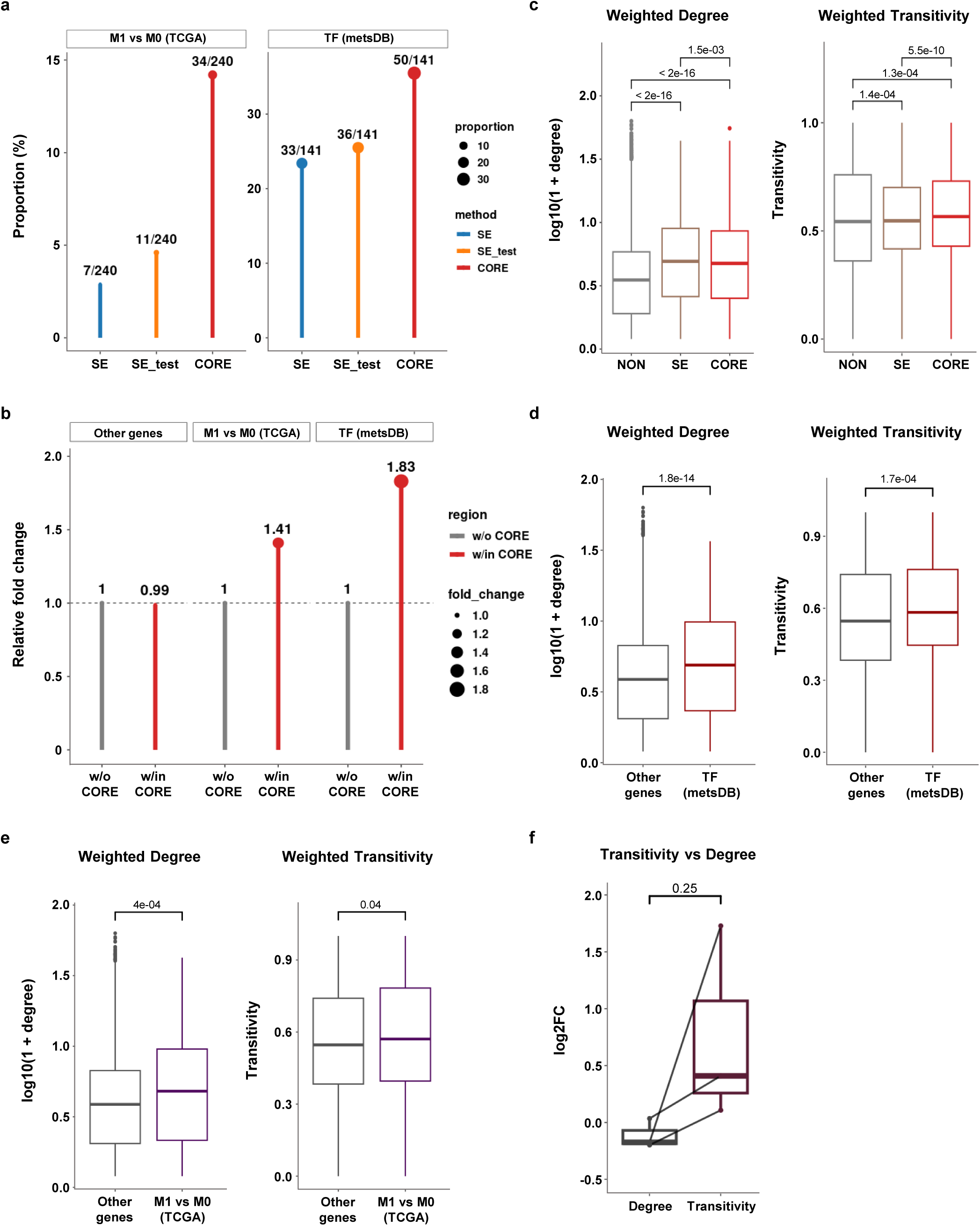
Metastasis-related programs preferentially reside within CORE-associated genes and link to distinct node-level network properties a, Proportion of CORE- and SE-associated genes that overlap with metastasis-related genes, relative to the total number of metastasis-related genes. M1 vs M0 (TCGA): differentially expressed genes between M1 (metastasis) and M0 (no metastasis) in TCGA GDC RNA-seq data, TF (metsDB): differential TFs between metastatic tumor and primary CRC based on TF regulon activity in scRNA-seq data. **b,** Relative fold changes in the number of metastasis-related genes overlapping genes annotated from peaks within or without COREs. Other genes: genes associated with H3K4me1 peaks that do not overlap two categories, M1 vs M0 (TCGA) and TF (metsDB). **c,** Box plots showing weighted degree and weighted transitivity (weighted local clustering coefficients) in CORE, SE, and NON. CORE: H3K4me1 peaks within COREs, SE: H3K4me1 peaks within SEs, NON: H3K4me1 peaks that do not fall into either COREs or SEs. Correlation coefficients are used as edge weights in the co-accessibility network. **d,** Box plots showing weighted degree and weighted transitivity (weighted local clustering coefficients) in peaks associated with TF (metsDB) and Other genes. **e,** Box plots showing weighted degree and weighted transitivity (weighted local clustering coefficients) in peaks associated with M1 vs M0 (TCGA) and Other genes. **f,** Log2-transformed fold changes of mean weighted degree and mean clustering coefficient between COREs and SEs. P-values were calculated to measure statistical significance (**c**, Kruskal-Wallis test with Dunn’s test. P-values were adjusted by the Holm correction method; **d,e**, two-sided Wilcoxon rank sum test; **f,** two-sided Wilcoxon signed-rank test).

**Extended Data Fig.10:**
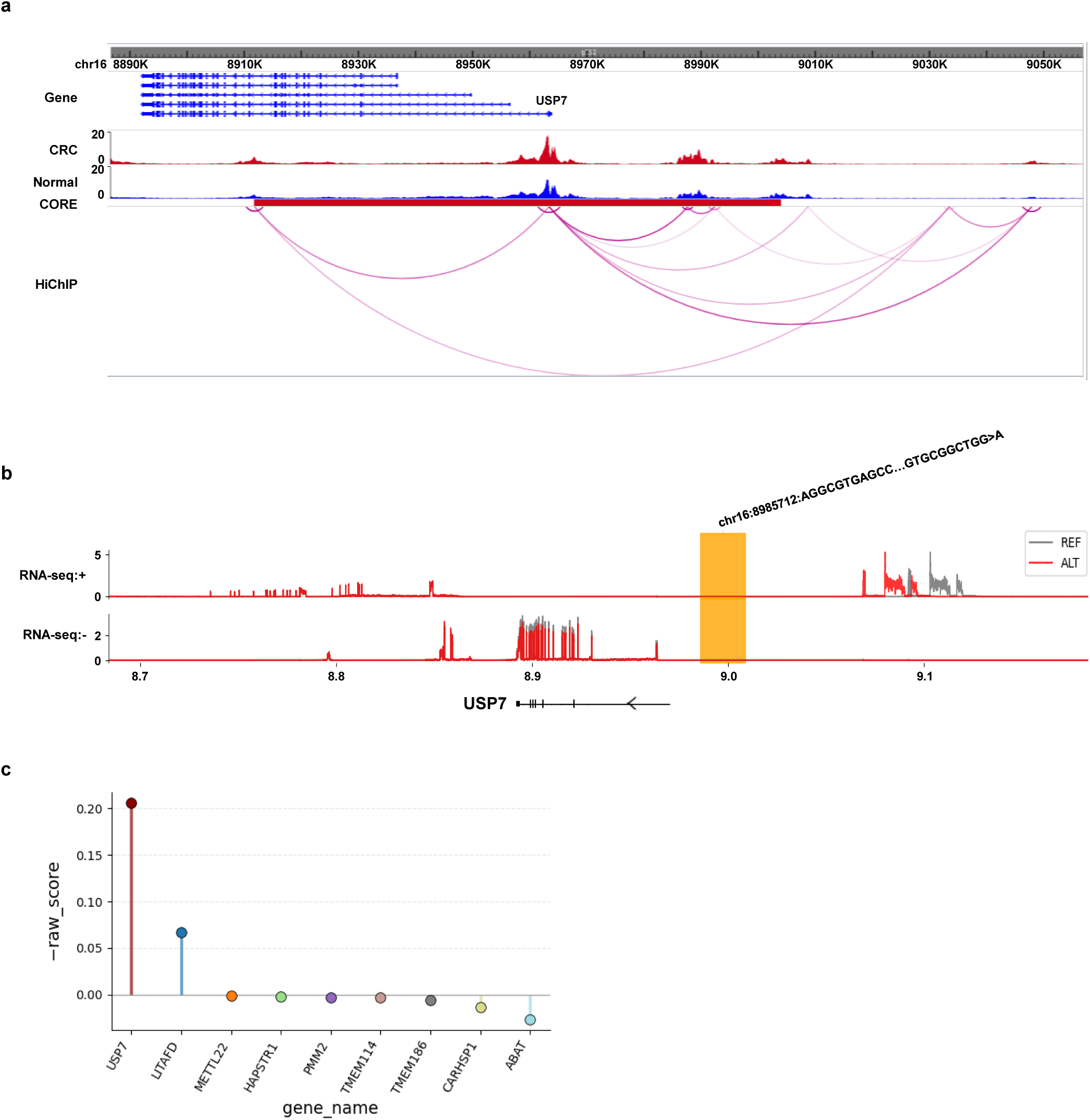
***In silico* ablation of a USP7-proximal CORE constituent enhancer predicts reduced transcriptional output a,** Track visualization at the *USP7* locus with H3K27ac HiChIP interactions from HT29 cells and normalized H3K27ac ChIP-seq signals in CRC and Normal samples. Colored and annotated by disease state: CRC (red color), Normal (blue color). **b,** Track visualization of RNA-seq from the Left Colon ontology under *in silico* knockout of the CORE-associated enhancer in the vicinity of *USP7*. **c,** Magnitude of the predicted effect on RNA-seq reads from the Left Colon ontology under the same perturbation.

**Supplementary Fig.1:**
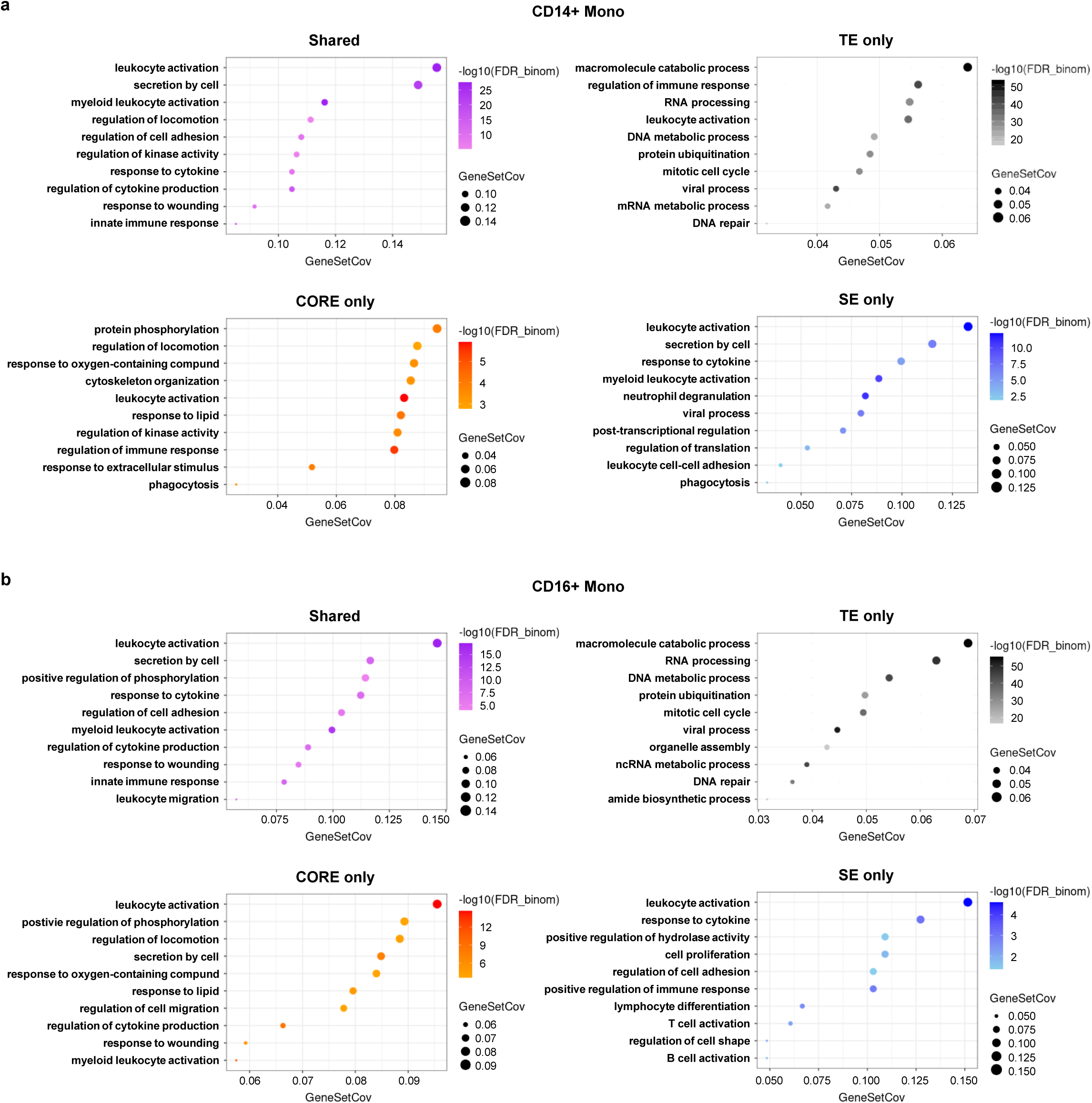
GO enrichment analysis of CORE in monocyte subpopulation a, GO enrichment analysis in CD14+ Monocytes. **b,** GO enrichment analysis in CD16+ Monocytes. Shared: Regions shared between COREs and SEs, CORE only: Regions exclusively found in COREs, SE only: Regions exclusively found in SEs, TE only: Regions exclusively found in TEs. Dots are colored by –log10(binomial FDR-adjusted P-value), and the dot sizes indicate the Gene Set Coverage.

**Supplementary Fig.2:**
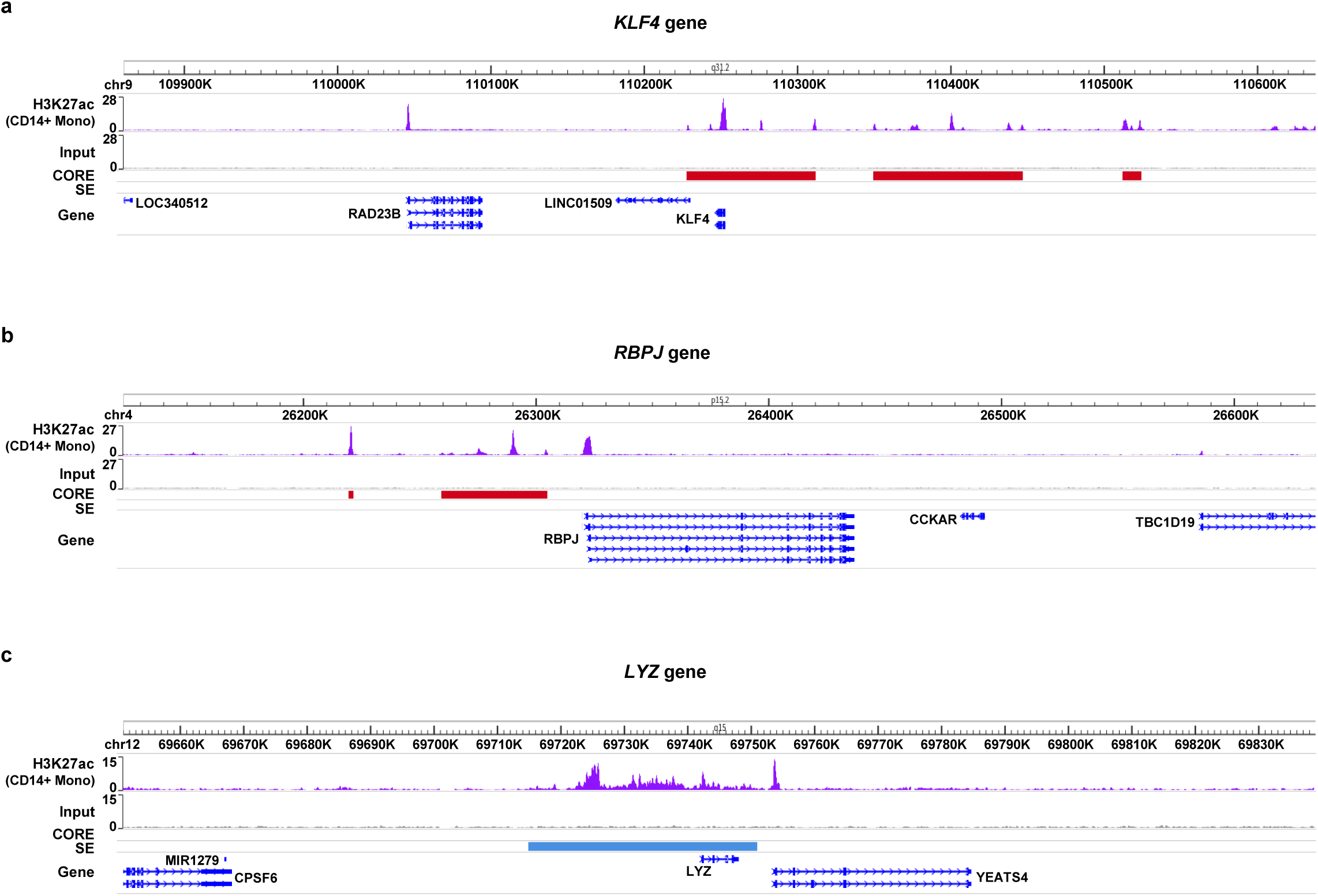
Track visualization of KLF4, RBPJ, and LYZ in CD14+ monocytes a, Track visualization at the *KLF4* locus with normalized H3K27ac ChIP-seq signals. **b,** Track visualization at the *RBPJ* locus with normalized H3K27ac ChIP-seq signals. **c,** Track visualization at the *LYZ* locus with normalized H3K27ac ChIP-seq signals. Colored and annotated by region categories: COREs (red color), SEs (blue color).

**Supplementary Fig.3:**
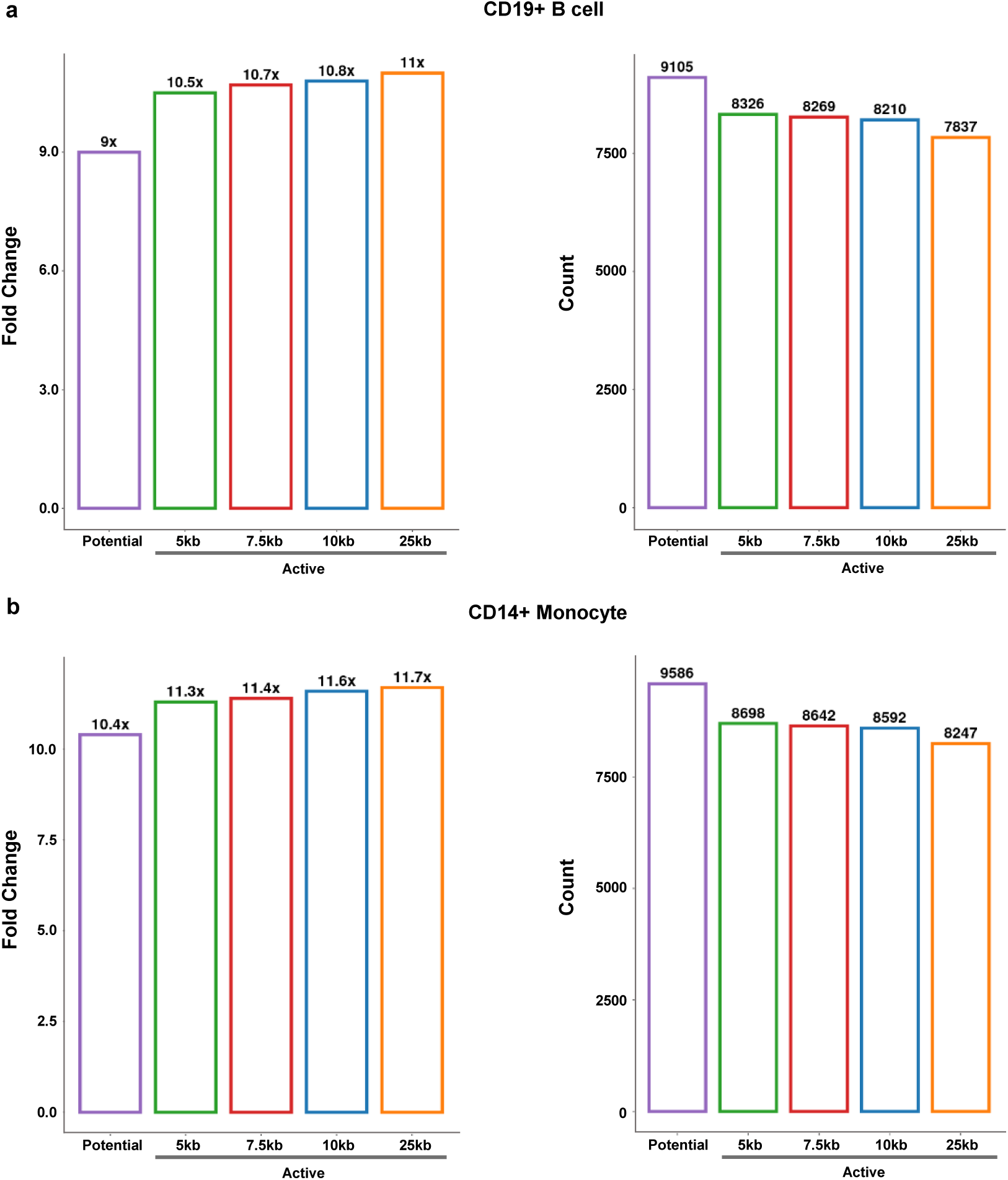
Active-option CORE shows a higher active-to-poised enhancer ratio than potential-option CORE a, Bar plots showing the fold change and the number of active enhancers in CD19+ B cells across different CORE calling options. **b,** Bar plots showing the fold change and the number of active enhancers in CD14+ Monocytes across different CORE calling options. Left: fold change in the number of active enhancers relative to the number of poised enhancers, Right: The number of active enhancers. Borderlines are colored by CORE calling options: Potential (COREs from the potential option; purple color), 5kb (COREs from the active option using 5kb as proximal enhancer distance; green color), 7.5kb (COREs from the active option using 7.5kb as proximal enhancer distance; red color), 10kb (COREs from the active option using 10kb as proximal enhancer distance; blue color), 25kb (COREs from the active option using 25kb as proximal enhancer distance; orange color).

**Supplementary Fig.4:**
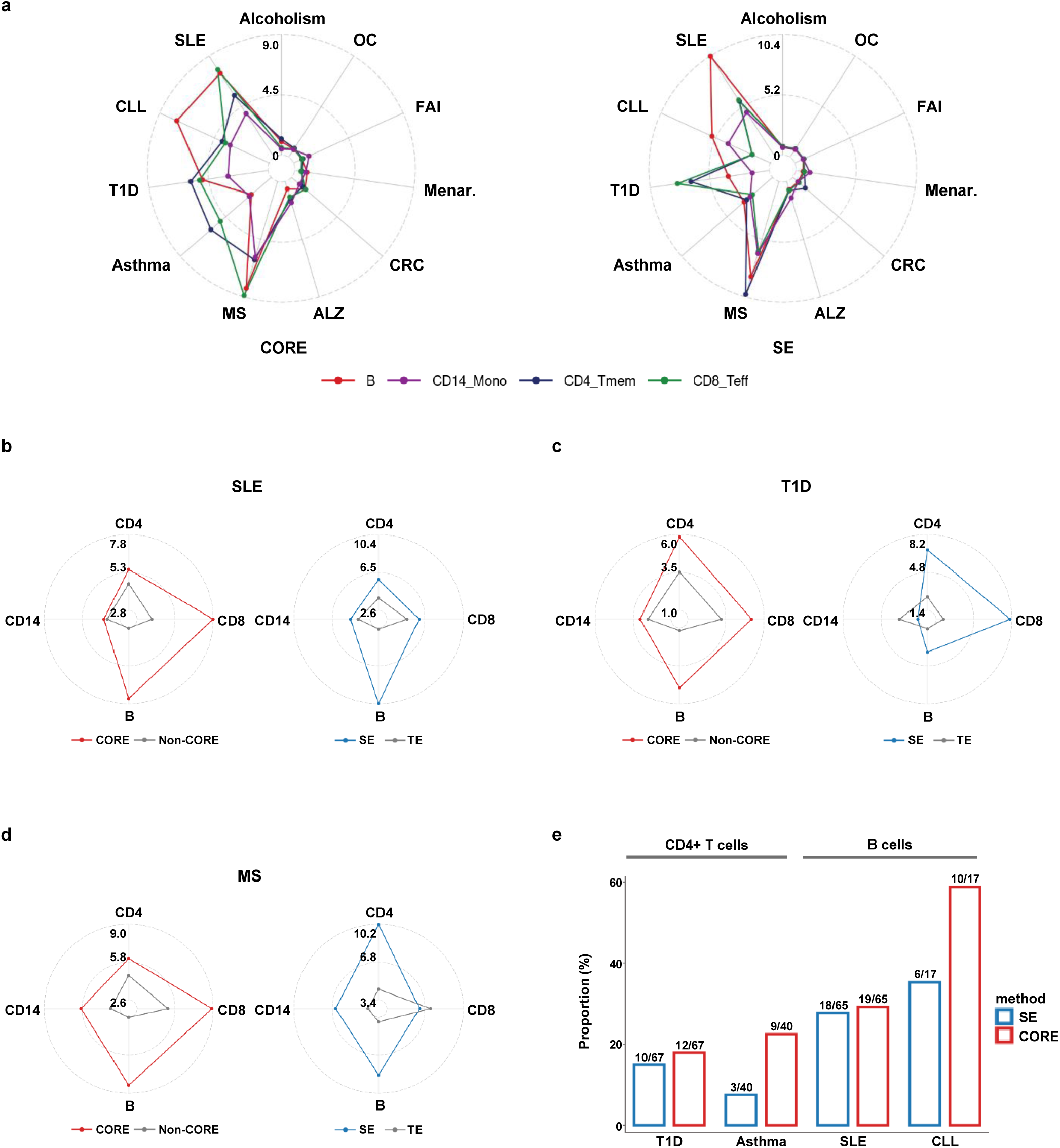
Immune-related disease GWAS variants (Hnisz *et al.*) are enriched in COREs a, Enrichment analysis of GWAS variants for immune-related diseases or non-immune-related traits in COREs (left) and SEs (right). SLE: Systemic Lupus Erythematosus, CLL: Chronic Lymphocytic Leukemia, T1D: Type 1 Diabetes, Asthma: Asthma, MS: Multiple Sclerosis, ALZ: Alzheimer’s disease, CRC: Colorectal Cancer, Menar.: Menarche, FAI: Fasting Insulin, OC: Ovarian Cancer, Alcoholism: Alcoholism. **b-d,** Enrichment analysis of GWAS variants for **b,** SLE; **c,** T1D; **d,** MS in COREs and SEs. COREs (red color), SEs (blue color), and TEs (grey color). B: B cells, CD14: CD14+ Monocytes, CD4: CD4+ Memory T cells, CD8: CD8+ Effector T cells. **e,** Proportion of GWAS variants that overlap with COREs or SEs, relative to the total number of GWAS variants in each autoimmune disease category. Borderlines are colored by method: COREs (red color), SEs (blue color).

**Supplementary Fig.5:**
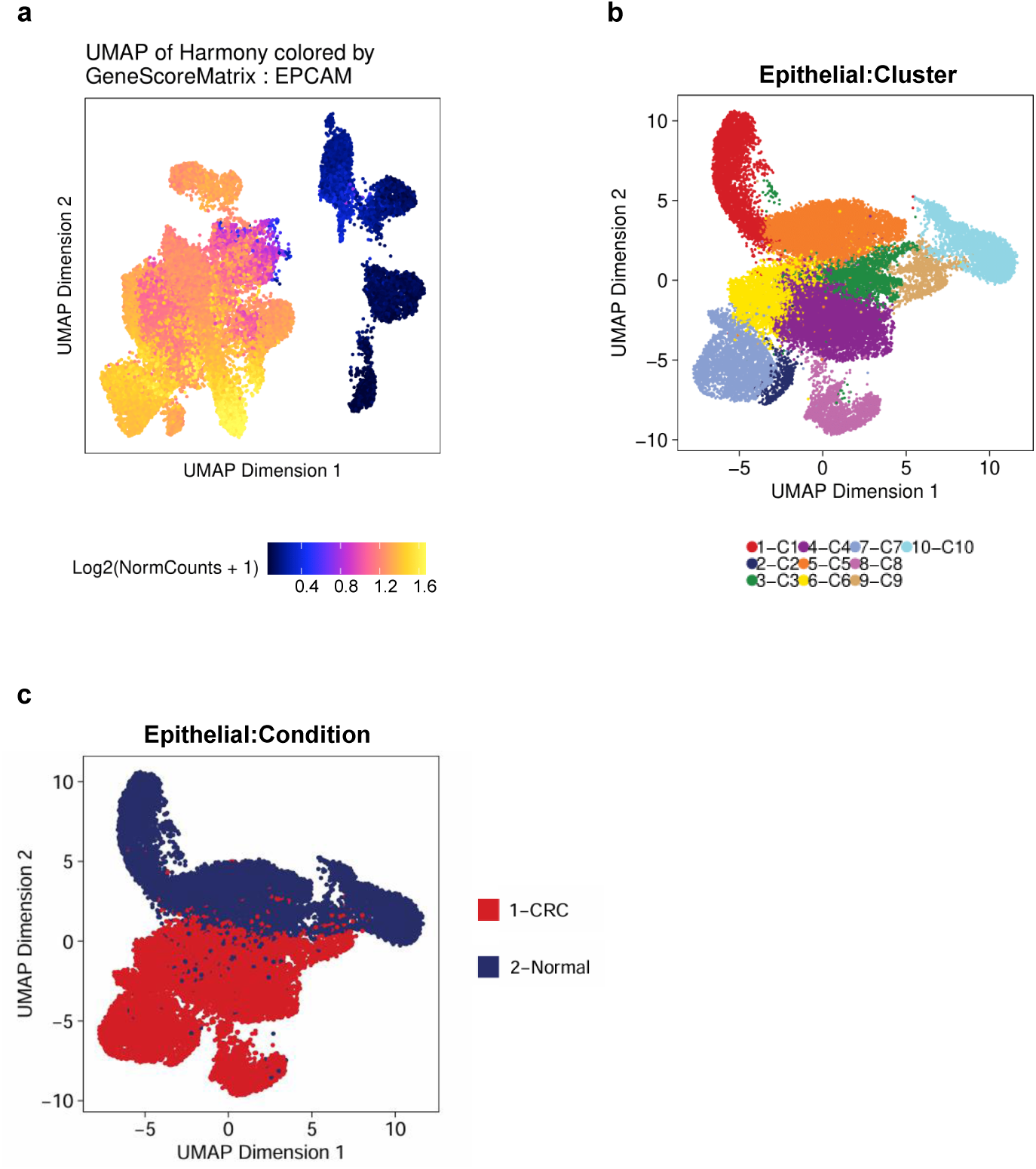
Characterization of CRC epithelial cell clusters and canonical marker expression a, UMAP embedding of the CRC scATAC-seq dataset colored by *EPCAM* gene expression. **b,** UMAP embedding of the CRC scATAC-seq dataset colored by sub-cluster in epithelial cells. **c,** UMAP embedding of the CRC scATAC-seq dataset colored by condition (disease state) information. CRC (Colorectal cancer epithelial cells; red color), Normal (Normal epithelial cells; blue color).

**Supplementary Fig.6:**
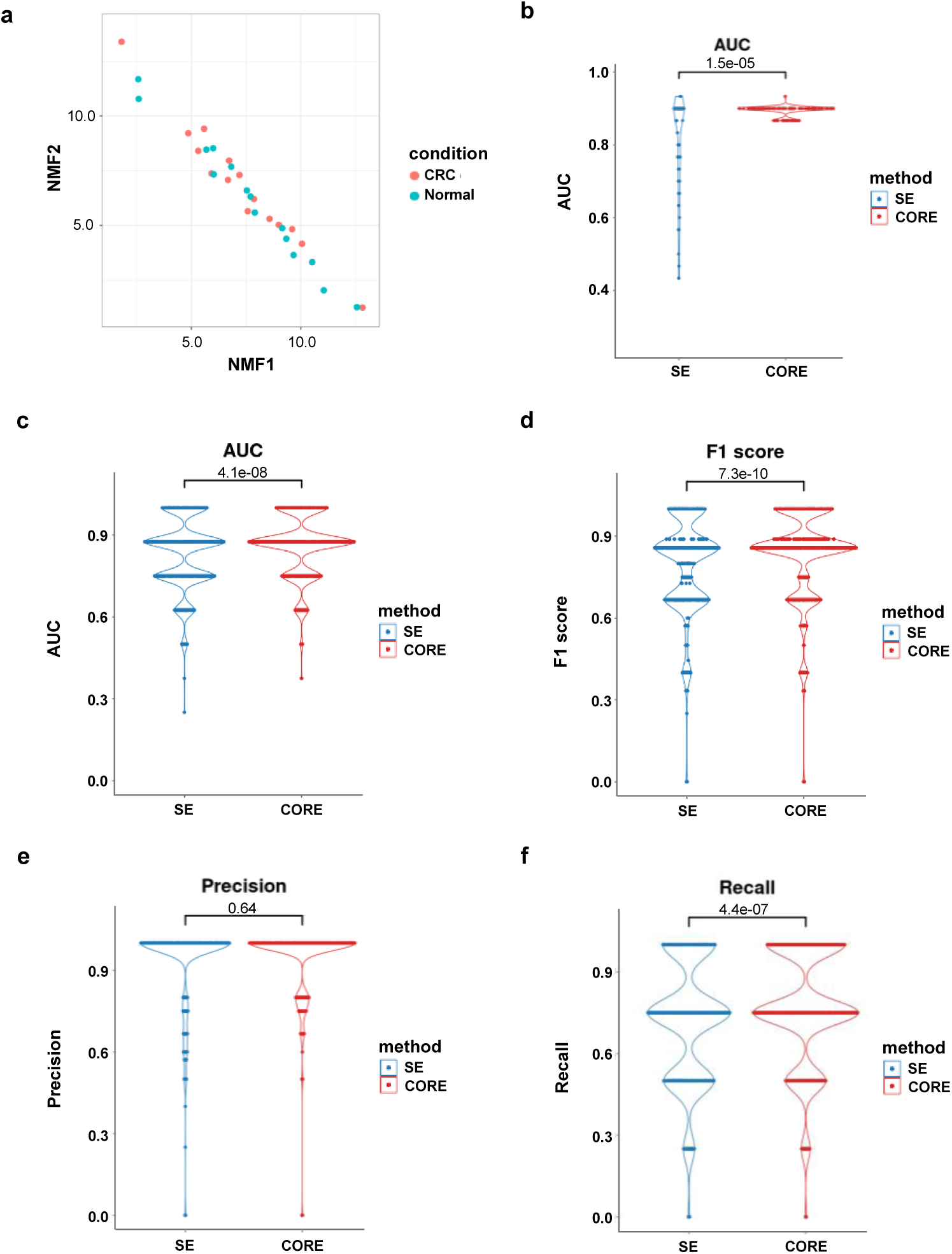
Comparison of classification performance between COREs and SEs (GSE156613) a, NMF analysis of H3K27ac ChIP-seq signals within SEs to distinguish between CRC and Normal samples. Dots are colored by disease state: CRC (red color), Normal (blue color). **b,** Violin plot showing classification performance between COREs and SEs. Area Under Curve (AUC) values were calculated by K-means clustering on the NMF-reduced dimension. **c-f,** Violin plot showing XGBoost classification performance between COREs and SEs. **c,** AUC (Top Left), **d,** F1 score (Top Right), **e,** Precision (Bottom Left), and **f,** Recall (Bottom Right). P-values were calculated to measure statistical significance (**b-f**, two-sided Wilcoxon rank sum test).

**Supplementary Fig.7:**
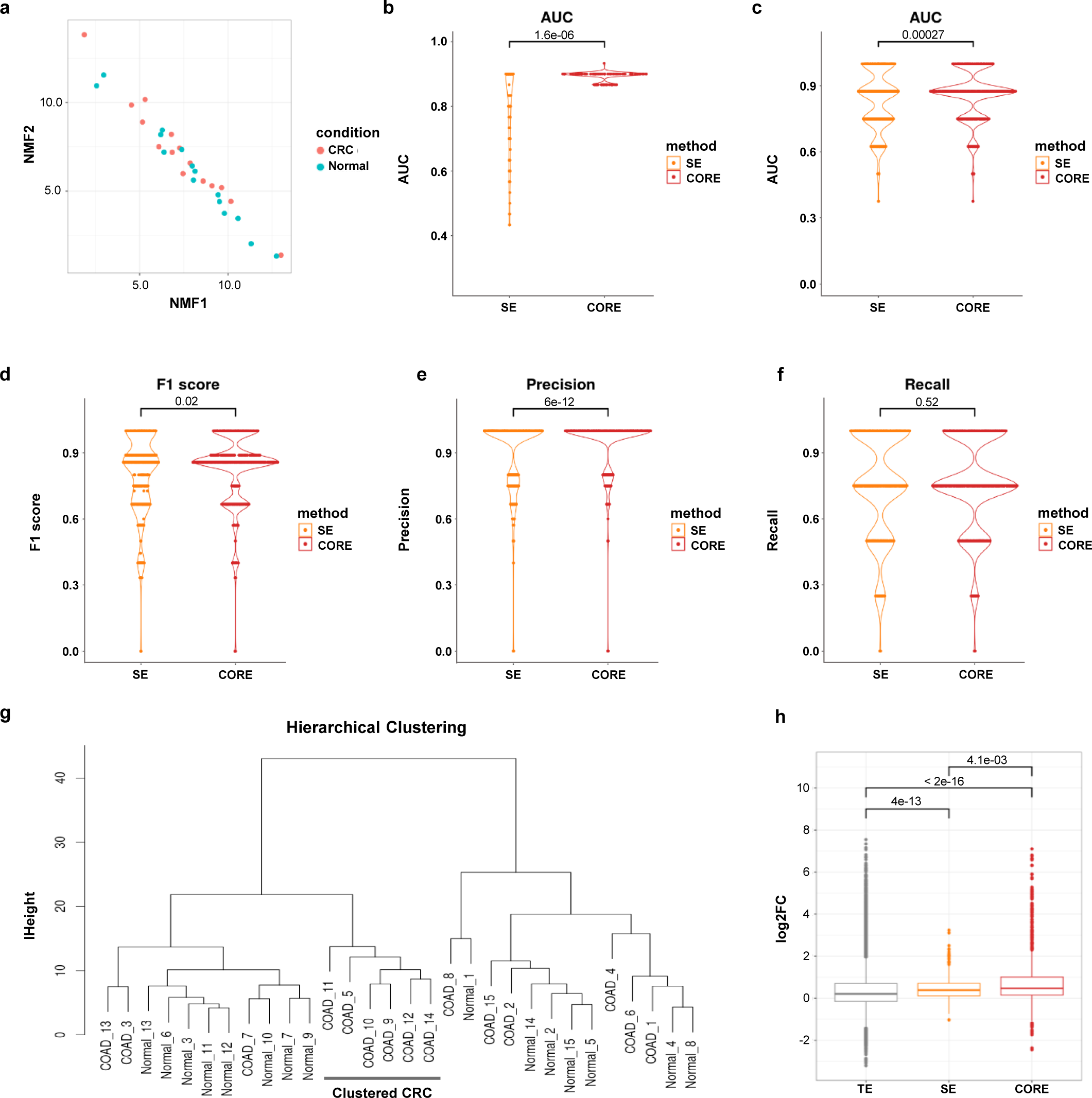
Comparison of classification performance between COREs and SEs (GSE136889) a, NMF analysis of H3K27ac ChIP-seq signals within SEs to distinguish between CRC and Normal samples. Dots are colored by disease state: CRC (red color), Normal (blue color). **b,** Violin plot showing classification performance between COREs and SEs. Area Under Curve (AUC) values were calculated by K-means clustering on the NMF-reduced dimension. **c-f,** Violin plot showing XGBoost classification performance between COREs and SEs. **c,** AUC (Top Right), **d,** F1 score (Bottom Left), **e,** Precision (Bottom Middle), and **f,** Recall (Bottom Right). **g,** Hierarchical clustering result for CRC and Normal samples. Euclidean distance was used as a distance metric for clustering. The cluster containing the most similar CRC samples identified using SEs was labeled as Clustered CRC. **h,** Log2-transformed fold changes of H3K27ac ChIP-seq signals between CRC and Normal within COREs, SEs, and TEs. Dots and borderlines are colored by region categories: COREs (red color), SEs (orange color), TEs (grey color). P-values were calculated to measure statistical significance (**b-f**, two-sided Wilcoxon rank sum test; **h**, Kruskal-Wallis test with Dunn’s test. P-values were adjusted by the Holm correction method).

**Supplementary Fig.8:**
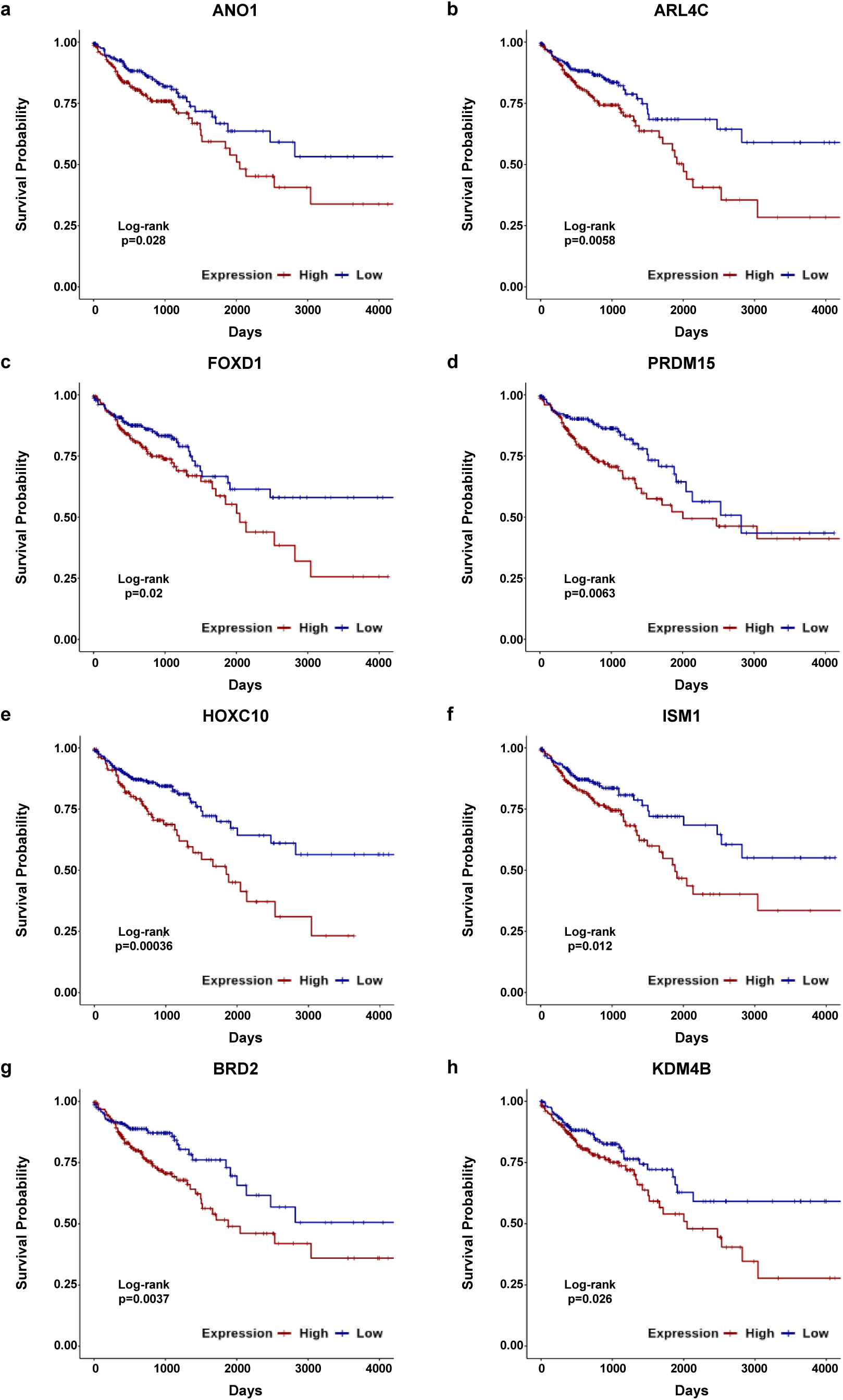
Survival analysis for CORE-associated genes a-h, Kaplan-Meier survival analysis of CORE-associated genes in TCGA CRC samples. **a,** *ANO1*; **b,** *ARL4C*; **c,** *FOXD1*; **d,** *PRDM15*; **e,** *HOXC10*; **f,** *ISM1*; **g,** *BRD2*; **h,** *KDM4B*. Lines are colored by expression level status: High gene expression (red color), Low gene expression (blue color). Survival curves were obtained from RNA-seq data. P-values were calculated using the Log-rank test.

**Supplementary Fig.9:**
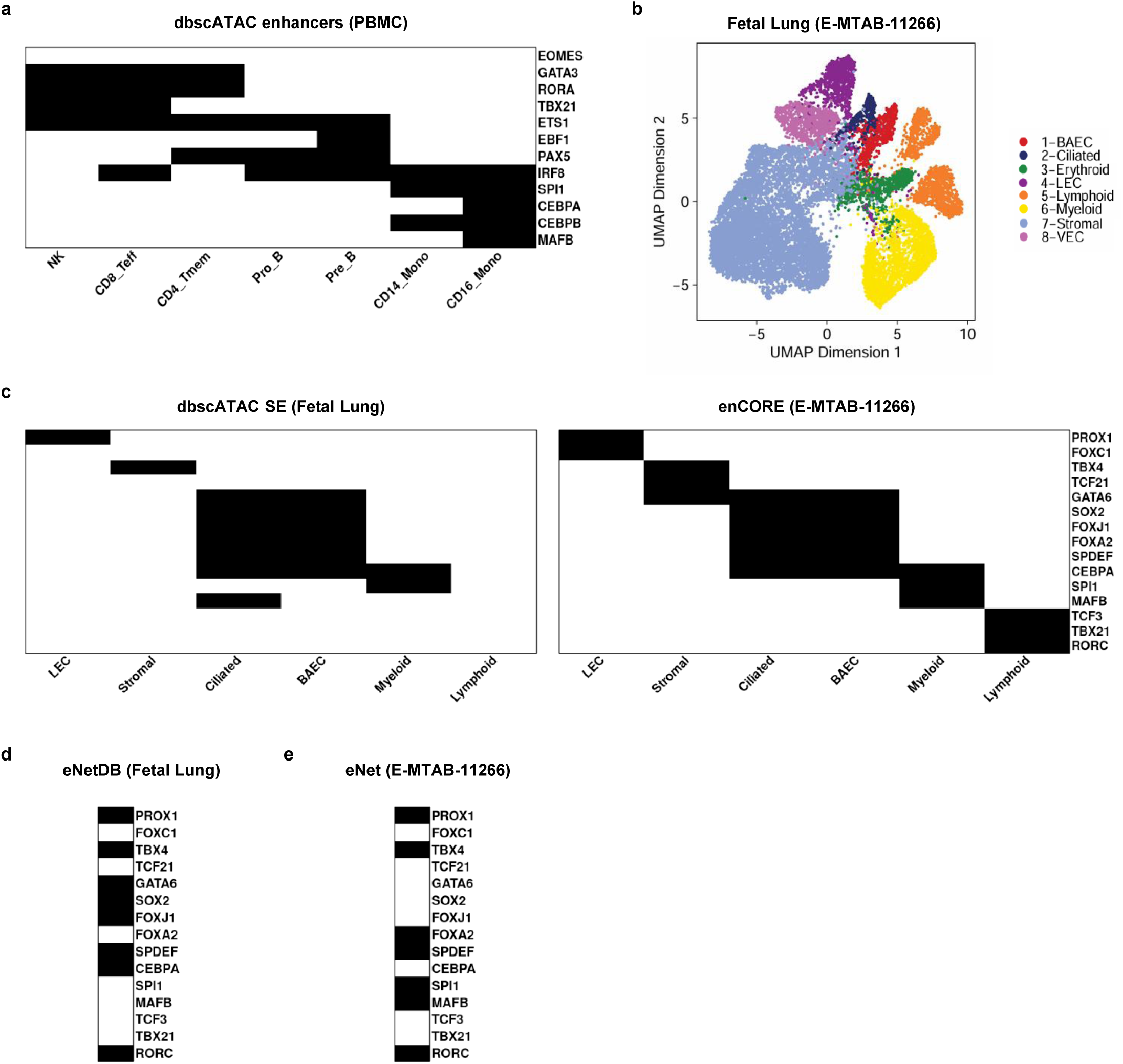
enCORE supports robust recovery of cell-type-specific master transcription factors under single-cell sparsity a, Heatmap for cell type-specific master regulators in PBMC. Black: gene present in the dbscATAC PBMC single cell enhancer-associated genes of the given cell type. White: absent. **b,** UMAP embedding of the Fetal Lung scATAC-seq dataset (E-MTAB-11266), colored and annotated by cell type. **c,** Heatmap for cell type-specific master regulators in fetal lung. Region category: SEs derived from dbscATAC human fetal lung (left) and COREs derived from E-MTAB-11266 (right). Black: gene present in the region category-associated genes of the given cell type. White: absent. **d,** Heatmap for cell type-specific master regulators in fetal lung. Black: gene present in eNet hub genes derived from eNetDB. White: absent. **e,** Heatmap for cell type-specific master regulators. Black: gene present in the eNet hub genes derived from E-MTAB-11266. White: absent.

**Supplementary Fig.10:**
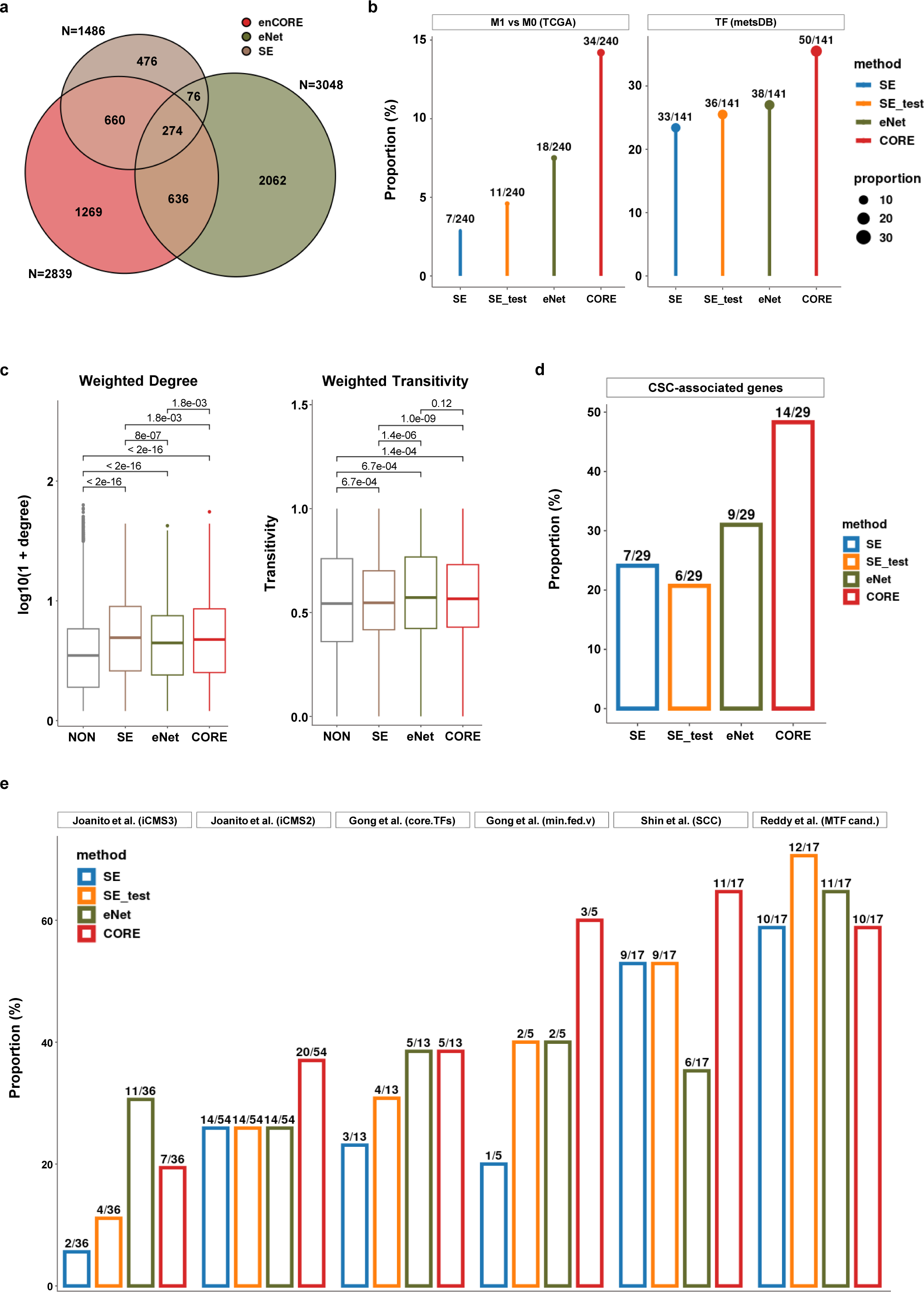
enCORE identifies a topology-defined enhancer profile distinct from SE and eNet in CRC a, Euler diagram showing the overlap among CORE/SE-associated genes and eNet hub genes. **b,** Proportion of CORE/SE-associated genes and eNet hub genes that overlap with metastasis-related genes, relative to the total number of metastasis-related genes. M1 vs M0 (TCGA): differentially expressed genes between M1 (metastasis) and M0 (no metastasis) in TCGA GDC RNA-seq data, TF (metsDB): differential TFs between metastatic tumor and primary CRC based on TF regulon activity in scRNA-seq data. **c,** Box plots showing weighted degree and weighted transitivity (weighted local clustering coefficients) in CORE, SE, eNet, and NON. CORE: H3K4me1 peaks within COREs, SE: H3K4me1 peaks within SEs, eNet: H3K4me1 peaks that overlap eNet hub enhancers, NON: H3K4me1 peaks that do not overlap COREs, SEs, or eNet hub enhancers. Correlation coefficients are used as edge weights in the co-accessibility network. Kruskal-Wallis test with Dunn’s test. P-values were adjusted by the Holm correction method. **d,** Proportion of CORE/SE-associated genes and eNet hub genes that overlap with cancer stem cell (CSC)-associated genes, relative to the total number of CSC-associated genes. **e,** Proportion of CRC master TF candidates that overlap genes associated with each region category, relative to the total number of CRC master TF candidates in each study. Region category: COREs (CORE; red color), eNet (eNet; green color), SEs from GSE136889 (SE_test; orange color), and SEs from GSE156613 (SE; blue color).

## Notes

### Competing Interest Statement

The authors have declared no competing interest.

### Summary of Updates

Major Changes 1. Corrected a potentially misleading typo and the accompanying description in the Methods definition of the Specificity score for cell type X (nTPM_{X^C,gene_{X^C}}). Minor Changes 1. Replaced "autoimmune" with "immune-related" in the Abstract, as the representative examples (CLL, Asthma, and CAD) are immune-related diseases but not autoimmune diseases. 2. Made several minor, non-substantive edits throughout the main text and Methods (e.g., removing unnecessary statements and adjusting wording) that do not affect the overall conclusions.

